# Beyond statistical significance: ranking transcription factor binding motifs by effect size

**DOI:** 10.64898/2026.06.19.732679

**Authors:** Coby Viner, Scott Mastromatteo, Danielle Denisko, Jeffrey Negrea, Yanbo Tang, Lin Zhang, Michael M. Hoffman, Lei Sun

## Abstract

Chromatin immunoprecipitation-sequencing (ChIP-seq) has wide use in identifying transcription factor binding sites. DNA sequence motifs specific to a targeted transcription factor occur more frequently near ChIP-seq peak centres. The most common approach to quantifying relative motif enrichment ranks motifs by p-value. Because sample sizes can vary substantially across examined motifs, p-value magnitudes may reflect this heterogeneity rather than the biological effect of interest. As alternatives, we considered four ranking methods based on effect sizes: (a) a modified Cliff’s delta, (b) the lower bound of a frequentist asymptotic confidence interval, (c) the lower bound of a frequentist finite-sample confidence interval, and (d) the lower bound of a Bayesian credible region. Through extensive simulations, the four alternatives better recovered the simulated central-enrichment ordering under heterogeneous sample sizes. Using published ChIP-seq data for GATA3, the effect size methods ranked the known targeted motif highest, even compared to highly similar motifs for other GATA family members, while p-value ranking did not. In a separate SRF application, all four alternative methods also consistently ranked the known motif highest. We recommend the asymptotic confidence interval lower bound for its simplicity, ease of implementation, and intuitive interpretation. The software is freely available (https://github.com/ScottMastro/motif-ranking).

## Introduction

Transcription factors are proteins that regulate gene expression via recognition and binding to specific DNA motifs^45^. Assays such as chromatin immunoprecipitation-sequencing (ChIP-seq)^29,51^ identify genome-wide transcription factor binding sites as peak sets. Each peak marks a genomic sequence centred around a candidate binding event.

Given a set of peaks and a collection of candidate motifs, motif analysis seeks peak set-motif correspondence. A commonly used criterion for this correspondence consists in central enrichment of best motif matches within peaks. For a given ChIP-seq experiment targeting one transcription factor, our goal is to rank all candidate motifs by their correspondence to the peak set and to identify the motif most likely bound by the targeted factor.

ChIP-seq and similar assays use antibodies to pinpoint genomic locations of DNA-binding proteins. Multiple controls for these assays can include input, mock immunoprecipitation, or ideally knockout (KO) experiments. Input controls involve isolation of cross-linked and fragmented DNA without adding an antibody. Mock immunoprecipitation uses a non-specific antibody during the affinity purification step instead of the target antibody. KO experiments use mutations directed to the gene encoding the target transcription factor, yielding no expression, prior to immunoprecipitation. Prior work in our group investigated how experimental controls can improve motif identification. We previously explored differential analyses of these controls, and developed PeaKO to optimize p-value rankings^19^.

Computational tools such as Multiple EM for Motif Elicitation (MEME)^3,4^ or Discriminative Regular Expression Motif Elicitation (DREME)^2^ identify enriched sequence motifs within a set of target sequences. In ChIP-seq results, these sequence motifs tend to occur more frequently near the centre of peaks, regions of the genome with high sequence read coverage^6^.

Accurate identification of transcription factor binding sites poses great difficulties^22^, with experimental noise adding additional complexity^30^. Antibody quality, sequencing biases, and batch effects can all lead to a failure to recover sufficient peaks for accurate motif elucidation or can enrich for additional off-target peaks.

Biological confounders compound these technical issues. Elucidation of the target motif identity proves especially problematic if the antibody has a slight affinity for other proteins of the same family^34^, as it may enrich motifs that are highly similar to the target. Additionally, certain classes of transcription factor binding profiles, referred to as “zingers”^66^, tend to systematically appear in many datasets, regardless of the antibody employed. All these factors interfere with our ability to directly detect motifs of interest.

Because direct identification of the target motif is often inconclusive, a common alternative is to rank candidate motifs by their central enrichment within ChIP-seq peaks. Here, we examine how such rankings are affected by motif-specific sample-size heterogeneity. We show that ranking by p-value can prioritize motifs with a larger number of contributing sequences. By contrast, ranking by effect size more directly quantifies the magnitude of central enrichment. Therefore, we consider and compare four alternatives based on effect size to identify a practical approach that is easy to implement and easy to interpret.

### CentriMo ranks motifs by enrichment in peak centres

The most common approach to quantitatively rank motifs enriched in peak centres uses p-values. For example, CentriMo^6^, part of the MEME Suite^7,8^, is a commonly used local motif enrichment analysis tool, which can determine the likely direct DNA-binding sequence motifs. CentriMo requires a predefined set of candidate motifs (here, from the JASPAR CORE 2016 vertebrates collection^42^, supplemented by *de novo* motifs discovered from the ChIP-seq data; see Methods).

To clarify the terminology used throughout: each ChIP-seq peak yields one input sequence of fixed length *ℓ*, extracted from the genome and centred on the peak summit. For each candidate motif, CentriMo scans all input sequences and records the position of the best match. Only sequences containing a match for a given motif contribute to that motif’s analysis; the number of contributing sequences defines the per-motif sample size *n*. Because different motifs match different subsets of sequences, *n* varies substantially across motifs within a single experiment—from dozens to tens of thousands in the datasets we examine. This variation in *n* is the source of the sample-size heterogeneity that motivates our work.

CentriMo then ranks the motifs under the assumption that biologically relevant motifs are more likely centrally aligned or enriched near the centre of the ChIP-seq peaks, using p-values that quantify the significance of central enrichment, as described below.

To obtain the p-value for each motif, CentriMo^6^ employs a one-tailed binomial test for central enrichment. Given a set of *n* input sequences of equal length *ℓ*, CentriMo defines the centre of each sequence to be position 0. A central window of length *W*, where *W* < *ℓ*, is then defined over exactly *W* position offsets relative to the centre. If we denote the start positions as integers, the central window starts at −⌊*W*/2⌋ and ends at ⌈*W*/2⌉ − 1, inclusive. For example, if *W* = 4, the central window includes positions {−2, −1, 0, 1}. For each of the *n* input sequences, the position of the best motif match may land either within or outside this central window. For simplicity, window membership is determined by the starting position of the best match, not the full span of the motif. Across all sequences, the number of matches within the central window, *X*, is assumed to follow a binomial distribution, with parameters *n* and *θ* ∈ [0, 1], *X* ∼ B(*n, θ*). Under the null hypothesis of no central enrichment, the best motif match should occur uniformly across the *ℓ* − *d* + 1 possible motif positions, where *d* is the width of the motif. Thus, under the null hypothesis *H*_0_, the probability, *θ*_0_, of observing the best motif match within the central window is *θ*_0_ = *W*/(*ℓ* − *d* + 1). Assuming independent sequences, the p-value is

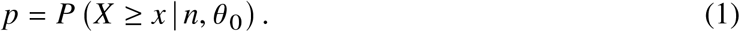

Finally, CentriMo ranks each motif based on its Bonferroni-corrected p-value.

### Ranking by p-value has limitations that can mislead

When ranking peak set-motif correspondence by p-value, motifs found in greater numbers of input sequences receive smaller p-values regardless of their enrichment magnitude. This problem is exacerbated by the highly heterogeneous sample sizes found for matches to distinct motifs in a single ChIP-seq experiment. The number of matches to a given motif can range from only 100 to more than 10 000. In these cases, the magnitude of p-values is dominated by this sample size heterogeneity, as opposed to the true effect size of interest^50,57,63^.

From a statistical perspective, this issue may seem unexpected, since p-value use in other genomic contexts is ubiquitous and generally accurate. Those other contexts, such as genome-wide association studies, have vastly less sample size heterogeneity^56^.

Furthermore, in ChIP-seq data most motifs are centrally located^6,65^, with p-values < 1 × 10^−50^, providing little discriminatory power. When nearly all p-values are astronomically small, the ranking among motifs depends on fine differences in p-value magnitude, which reflect sample-size variation rather than biological signal.

Consider an illustrative example of ranking two motifs based upon which one is more centrally enriched, relative to a mean-symmetric binomial distribution, under the null hypothesis (*θ*_0_ = 0.5). Suppose that for motif A, we have *n* = 100 input sequences and *X* = 80 matches within the central window. Thus the p-value for motif A, *p*_A_ = *P* (*X* ≥ 80 | *θ*_0_ = 0.5, *n* = 100) = 9.87 × 10^−10^. For motif B, suppose we have *n* = 1000 and *X* = 600. This instead yields *p*_B_ = *P* (*X* ≥ 600 | *θ*_0_ = 0.5, *n* = 1000) = 1.27 × 10^−10^, indicating that motif B has greater significance. This p-value, however, depends upon *n*, the number of sequences, and reflects the heterogeneity in sample sizes (100 versus 1000), rather than the difference in effect sizes. Specifically, the effect size—the departure of the observed proportion from its null value—is 80/100 − 0.5 = 0.30 for motif A, but only 600/1000 − 0.5 = 0.10 for motif B. Despite motif A having a threefold larger effect size, motif B receives the smaller p-value. Motif B’s more significant p-value is driven entirely by its tenfold larger sample size.

Given the inconsistent meaning of statistical significance in the face of such sample size heterogeneity, we focus on ranking motifs by effect size, using methods that separate the magnitude of central enrichment from the number of available sequences. More generally, the importance of effect size considerations has been widely highlighted^11,23,59^, motivating work to further demonstrate its importance in other domains.

### Effect sizes provide an alternative order for motif ranking

Here, we present several possible remedies to this problem, focusing on using effect sizes, instead of p-values, to address which motifs are more centrally enriched within their input sequences. Our methods include a modified Cliff’s delta^14,15^, a non-parametric method used for other genomics tasks^25,40^. We also utilize more standard approaches, including frequentist confidence intervals and Bayesian credible regions^27^. In total, we examine five ranking methods, including CentriMo p-values and four methods based on effect sizes (Table 1).

**Table 1.** Overview of the five ranking methods considered in this work. Generally, larger values of the ranking statistic indicate stronger central enrichment.

| Method | Ranking statistic | Inference type |
| --- | --- | --- |
| CentriMo p-value<br>Compares best-match positions to binomial distribution under null hypothesis $\theta_{i_0} = W_i/(\ell - d_i + 1)$<br>Contained in $[0, 1]$<br>Exceptionally, for this method, smaller values indicate stronger central enrichments<br>Highly sensitive to sample size heterogeneity | $p_i$ (Equation 1) | Frequentist significance test |
| Modified Cliff’s delta<br>Compares best-match positions to uniform reference using signed pairwise comparisons of distances from the peak centre<br>Contained in $[-1, 1]$ | $\delta_i^*$ (Equation 4) | Non-parametric effect size |
| Asymptotic confidence interval lower bound<br>Compares the estimated central-window proportion $\hat{\theta}_i$ to the null proportion $\theta_{i_0}$ using the central limit theorem approximation with Bonferroni adjustment across motifs<br>Bounded above by $1 - \theta_{i_0}$, with no guaranteed lower bound; can be negative | $L_i$ (Equation 8) | Frequentist effect size interval |
| Finite-sample confidence interval lower bound<br>Compares the estimated central-window proportion $\hat{\theta}_i$ to the null proportion $\theta_{i_0}$ using Hoeffding’s inequality simultaneous bound with finite-sample coverage guarantee<br>Usually more conservative than the asymptotic confidence interval lower bound<br>Bounded above by $1 - \theta_{i_0}$, with no guaranteed lower bound; can be negative | $\zeta_i$ (Equation 11) | Frequentist effect size interval |
| Bayesian credible lower bound<br>Compares the posterior distribution of $\theta_i$ to the null proportion $\theta_{i_0}$ using a beta-binomial model with Jeffreys prior<br>Contained in $[-\theta_{i_0}, 1 - \theta_{i_0}] \subseteq [-1, 1]$; can be negative<br>Underlying credible region for $\theta_i$ guaranteed to lie in $[0, 1]$, unlike the frequentist confidence interval<br>Asymptotically equivalent to the frequentist interval approach | $r_i$ (Equation 12) | Bayesian effect size interval |

Through both simulation and application studies, we show that rankings from these alternative methods are largely consistent with one another and, under heterogeneous sample sizes, better reflect the magnitude of central enrichment than rankings based on p-value alone. In practice, p-values can still be used to assess statistical evidence for central enrichment. Because many motifs in ChIP-seq data can have extremely small p-values, however, the key question is often not whether a motif is statistically enriched, but how strongly enriched it is relative to other motifs. Among the four effect size alternatives considered here, our goal is to identify a ranking statistic that is robust to sample-size heterogeneity, easy to implement, and straightforward to interpret.

## Methods

### Datasets

We analysed a total of 4 publicly available ChIP-seq experiment datasets (Table 2), selected as representative examples across diverse experimental conditions. Three have a KO (or knockout-equivalent) control, whereas NF-κB is non-differential; we include the latter to compare the ranking methods where no knockout is available. Knockout experiments provide ground truth for motif identity: the motif whose central enrichment disappears upon knockout of the target transcription factor gene corresponds to the directly bound target. Such paired wild-type/knockout datasets are comparatively rare, which constrains dataset selection. Of these 4, two (GATA3 and SRF) overlap with datasets from our prior PeaKO work^19^, originally collected by Krebs et al.^31^, while we selected the other two—ATF1 and the non-differential NF-κB dataset—from Gene Expression Omnibus (GEO)^20^. ATF1 was assayed via CRISPR epitope tagging ChIP-seq (CETCh-seq)^52^, a related ChIP-seq variant that incorporates FLAG epitope tagging to the tail of a target transcription factor.

**Table 2.** ChIP-seq (or CETCh-seq for ATF1) datasets examined, with associated GEO accession numbers. Table adapted from Denisko et al.^19^.

| Factor Identifier | Species | Reference | Number of replicates |  |  |
| --- | --- | --- | --- | --- | --- |
|  |  |  | Wild type | Knockout | Input |
| ATF1-FLAG GSE72082 | Human | Savic et al.<sup>52</sup> | 2 | 2<sup>a</sup> | 0 |
| GATA3 GSE20898 | Mouse | Wei et al.<sup>64</sup> | 1 | 1 | 0 |
| NF-κB GSE63736 | Human | de Oliveira et al.<sup>b,18</sup> | 2 | 0 | 2 |
| SRF 10.5281/zenodo.3405482 | Mouse | Sullivan et al.<sup>58</sup> | 1 | 1 | 0<sup>c</sup> |

<sup>a</sup> The provided CETCh-seq data provides the equivalent of a knockout. It comes from a modified cell line with FLAG-tagged ATF1. When using FLAG as the ChIP-seq analyte, the anti-FLAG antibody enriches ATF1-bound DNA only in the tagged cells. Unmodified cells lack the epitope, so the same assay yields no specific signal in them. These untagged cells therefore serve as a built-in knockout-equivalent, rather than requiring a separately sequenced control sample. We accordingly list this as the number of wild type (WT) replicates, despite having only two total samples.
<sup>b</sup> Non-differential; L1236 cells for the NF-κB pathway components NF-κB1/p50 and NF-κB2/p52, RelA/p65, and RelB (2 biological replicates, across these 4 sub-units).
<sup>c</sup> We excluded the available input dataset because it came from a different, older DNA sequencer and lacked quality scores. Therefore, we conducted this analysis without an input control.

We selected these datasets using several criteria. First, we sought datasets of interest for their particular assay or method, such as knock-out datasets previously analysed with PeaKO^19^. Second, we aimed for representative datasets across a range of transcription factors. Third, we selected datasets that demonstrated ranking differences from CentriMo’s usual measures in our preliminary analyses. These analyses are intended to be representative of the impact and typical performance of the statistical methods we consider. We did not endeavour to comprehensively evaluate all available datasets, and these analyses are sufficient to suggest the utility of the proposed methods across diverse datasets.

We accessed datasets through GEO, except for the SRF dataset^24^, available on Zenodo (`10.5281/zenodo.3405482`). The ATF1 and NF-κB experiments come from human tissue, while the other experiments come from mouse tissue. Only GATA3 and SRF overlap with datasets from our prior PeaKO work^19^; the NF-κB analysis here uses the distinct L1236 dataset from `GSE63736`^18^.

### Motifs

As in Denisko et al.^19^, we downloaded the collection of vertebrate motifs in MEME format^8^ from the JASPAR CORE 2016 motif database, which consists of curated position weight matrices (PWMs) derived from *in vivo* and *in vitro* methods^42^. We identified each canonical motif from the JASPAR collection as the motif corresponding to the target transcription factor. We provided motifs to CentriMo^6^ for central enrichment analyses.

### Pre-processing, alignment, and peak calling

Unless otherwise specified, default parameters were used for all software tools. We processed the ATF1-FLAG, GATA3, and SRF datasets similarly to those in Denisko et al.^19^, as summarized below. Before alignment, we trimmed adapter sequences with Trim Galore (version 0.4.1)^33^ which uses Cutadapt (version 1.8.3)^41^. We assessed sequencing data quality using FastQC (version 0.11.5)^1^. We used Picard’s `FixMateInformation` and `AddOrReplaceReadsGroups` (version 2.10.5)^13^ and Genome Analysis ToolKit (GATK)’s `PrintReads` (version 3.6)^43^ to prevent GATK errors. Then, we aligned reads with BWA `bwa-aln` (version 0.7.15)^35^, the recommended aligner within the BWA suite for reads shorter than 70 bp^36^, using Sambamba (version 0.6.6)^60^ for post-processing. Here, we use mouse assembly NCBI m37/mm9 for all murine analyses, and GRCh37/hg19 for data from human cells. Because paired wild-type and knockout samples are aligned to the same assembly, relative central enrichment of motifs is unaffected by assembly version. Next, we called peaks using MACS2 (version 2.0.10)^68^ with parameters `-q 0.05`.

Exceptionally, for the NF-κB datasets (`GSE63736`), we performed neither alignment nor peak calling. Instead, we used the provided union peak set for L1236 (`GSE63736_L1236_nfkb_peaks.txt.gz`), comprising regions bound by NF-κB1/p50, NF-κB2/p52, RelA/p65, or RelB, and used UCSC liftOver to convert those NCBI 36/hg18 peaks to GRCh37/hg19, keeping all human analyses on a common assembly.

### Motif analyses

We employed MEME-ChIP (version 4.11.4)^37,38^ from the MEME Suite^7,8^ for motif analysis. MEME-ChIP performs motif discovery with complementary algorithms MEME^3,4^ and DREME^2^, and motif enrichment with CentriMo^6^. Unlike with our prior work on PeaKO^19^, we used a simplified non-differential workflow, which just runs MEME-ChIP, in same manner as for other non-differential motif analyses of ours^62^.

We extended MACS2 narrowPeak regions equidistantly from peak centres to create a uniform set of 500 bp centred peaks^37^. Then, we extracted underlying genomic sequences using BEDTools (version 2.23.0)^49^ `slop` from a repeat-masked genome. We masked the genome with Tandem Repeats Finder (TRF) (version 4.09)^10^ with options `-h -m -ngs` and parameters `2 7 7 80 10 50 500` for mouse (as originally done^10^), and options `2 5 5 80 10 30 200` for human (as recommended^21^).

### Ranking by a modified Cliff’s delta

To rank motifs, first we propose the use of Cliff’s delta^14,15^. This measure quantifies how often values sampled from one probability distribution exceed values in another. This non-parametric metric is linearly related to the Mann-Whitney U statistic. Cliff’s delta also relates to the Probability of Superiority: the probability that the score of a randomly selected individual from one group will exceed that of a randomly selected individual from another^39^.

For two probability distributions *X* and *Y*, Cliff’s delta *δ* is the difference of the probability that values sampled from one *x* ∈ *X* exceed those from the other *y* ∈ *Y* :

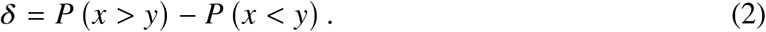

When used to compare observations from different populations, one can compute Cliff’s delta by comparing all pairwise combinations of values from these two populations *X* and *Y* . Considering each pair of values (*x* ∈ *X, y* ∈ *Y*), Cliff’s delta is the difference between the number of pairs where *x* > *y* and those where *x* < *y*, normalized by the total number of pairs. Cliff’s delta ignores any ties. The total number of pairs is the product of the cardinality of the populations |*X*||*Y* |.

We adapt Cliff’s delta (Equation 2) to compare motif correspondence to peak sets on the basis of central enrichment. For each pair (*x* ∈ *X, y* ∈ *Y*), we examine which value has the larger magnitude, |*x*| or |*y*|. Using magnitudes makes this comparison centre-oriented: it treats symmetrically those values at equal distance from the centre, regardless of sign. Specifically, we compute the sign function of the difference of the magnitudes, sgn (|*x*| − |*y*|), which is 1 if |*x*| is larger, −1 if |*x*| is smaller, and 0 if |*x*| = |*y*|. We then sum all of the sign functions and divide by the total number of pairs.

Cliff’s delta adapted for comparing central enrichment, therefore, is:

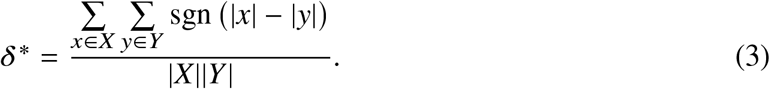

The resulting value lies within [−1, 1]. The superscript star emphasizes that this adaptation of the traditional Cliff’s delta compares magnitudes instead of potentially signed quantities.

In the context of peak set-motif correspondence, we search for *m* motifs across *n* peaks of equal length *ℓ*. For each peak-motif combination, we find the starting position of the best match. Motif *i* ∈ [1, *m*] has width *d*_*i*_. The multiset *V*_*i*_ contains the starting positions of the best motif matches. Let *ℓ*_*i*_ = *ℓ* − *d*_*i*_ + 1 denote the effective number of valid starting positions for motif *i*. We recentre these positions on the midpoint of motif *i*’s valid range, so that they lie within the interval [−*ℓ*_*i*_/2, *ℓ*_*i*_/2], whose length equals the number *ℓ*_*i*_ of valid starting positions. At position *j*, we identify the multiplicity *v*_*i,j*_ of best matches at *j*. Under the null hypothesis, we compare the observed distribution of *V*_*i*_ to a per-motif reference uniform distribution U([−*ℓ*_*i*_/2, *ℓ*_*i*_/2]).

Instead of randomly sampling from this uniform distribution, we take its reference population *U* to be the entire support interval [−*ℓ*_*i*_/2, *ℓ*_*i*_/2], whose length is |*U*| = *ℓ*_*i*_. Applying Equation 3 with *X* = *U* and *Y* = *V*_*i*_, in the continuous-population limit, yields a closed form for 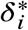:

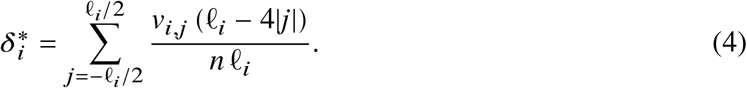

The linear weight (*ℓ*_*i*_−4|*j*|) is the measure of {*u* ∈ *U* : |*u*| > |*j*|} minus the measure of {*u* ∈ *U* : |*u*| < |*j*|}. This closed-form expression is equivalent to a previously-published sample estimate of Cliff’s delta^32^, as applied to motif central-enrichment ranking. One can rank motifs by values of 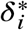. Greater values indicate motifs with greater central enrichment.

### Ranking by the lower bound of an asymptotic confidence interval

We now resort to the classical confidence interval approach^67^ to rank motifs. We rank motif *i* of width *d*_*i*_’s correspondence to a peak set with *n* input sequences under CentriMo’s^6^ assumptions. Specifically, we examine the number 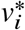 of best motif matches with starting positions that fall in the centre window of length *W*_*i*_:

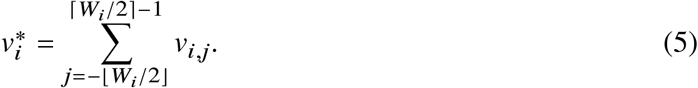

Window membership is determined by the starting position of the best match (not the full span of the motif), which is consistent with the null probability 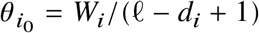 defined below.

We assume peak set-motif correspondence when the starting positions of the best motif matches 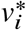 which fall in the centre window of length *W*_*i*_ follow a binomial distribution. That is, 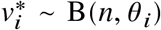. Under this assumption, we can estimate the binomial success parameter *θ*_*i*_ by dividing 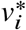 by the number *n* of peaks:

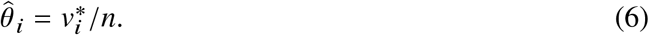

Under the null, 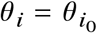 and 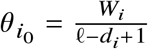. To rank the motifs by their estimated effect sizes, we define Δ_*i*_, the departure of the binomial success probability from its null value, 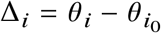. We can estimate 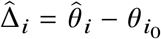. By the central limit theorem,

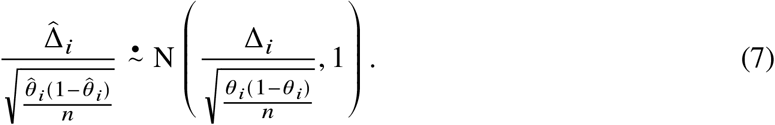

This follows from the central limit theorem for the binomial proportion: since 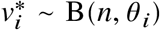, we have 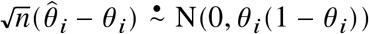. Subtracting Δ_*i*_ from 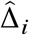, the constant 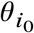 cancels: 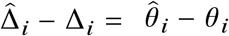. Therefore, the same asymptotic normality result holds: 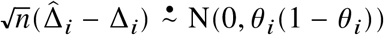. The variance parameter *θ*_*i*_(1 − *θ*_*i*_) gives a standard error of 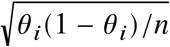 for 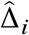; dividing by this standardizes to N(0, 1). Replacing the unknown *θ*_*i*_ with 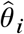 in the standard error (valid asymptotically by Slutsky’s theorem) gives the denominator 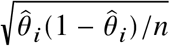 in the displayed result.

We will adjust confidence intervals using the conservative Bonferroni correction for multiple comparisons. Using the inverse cumulative distribution function of the standard normal distribution Φ^−1^, define 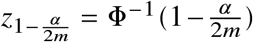 as its upper *α*/2*m* quantile. The factor of *m* in the denominator is the Bonferroni correction: to achieve simultaneous 100 (1 − *α*) % coverage—that is, probability at least 1 − *α* that every interval contains its true effect size—across all *m* motifs, each individual confidence interval is constructed at level 1 − *α*/*m*. The two-sided correction halves the tail probability to *α*/2*m*.

For a Type I error rate *α*, we can construct a 100 (1 − *α*) % asymptotic confidence interval for Δ_*i*_ for each motif *i*. The lower bound, *L*_*i*_, after the Bonferroni correction, is

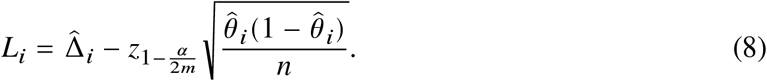

We can then use *L*_*i*_ to rank the motifs by the lower bounds of their confidence intervals.

### Ranking by the lower bound of a finite-sample confidence interval

We can also construct an alternative non-asymptotic confidence interval approach. This approach uses Hoeffding’s inequality^26^ to bound 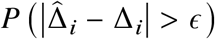 for any deviation threshold *ϵ* > 0. Under the same distributional assumption used above that 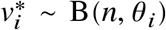, we may take 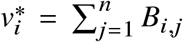 where 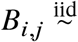 Bernoulli(*θ*_*i*_). Since 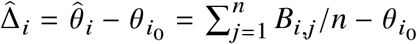 is an average of independent and identically distributed (i.i.d.) terms and each bounded within 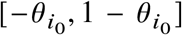, Hoeffding’s inequality states that

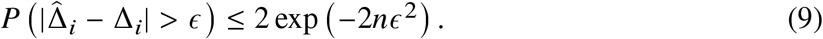

Setting 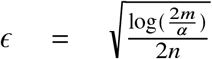 in the Hoeffding bound gives 2 exp(−2*nϵ*^2^) = *α*/*m*, so 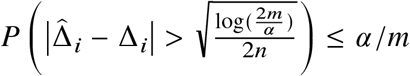 for each motif *i*. Applying the Bonferroni union bound across all *m* motifs and taking the complement then yields a simultaneous bound for all motifs:

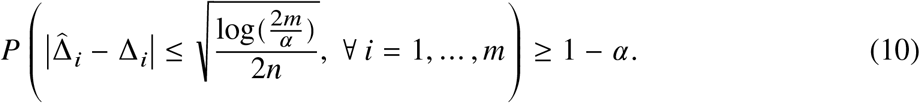

Similar to the asymptotic confidence interval approach, we can define the lower 100 (1 − *α*) % effect size of the *i*th motif as

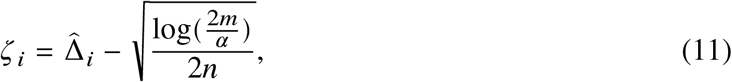

and use it to rank the motifs.

The differences between rankings based on *L*_*i*_ (Equation 8) and those based on *ζ*_*i*_ (Equation 11) will generally be small, when sample sizes are large and the binomial success probability *θ*_*i*_ is not close to 0 or 1. The finite-sample intervals from Hoeffding’s inequality are non-asymptotic, and therefore provide simultaneous finite-sample coverage under the stated assumptions.

In contrast, the asymptotic approach used for *L*_*i*_ relies on a normal approximation (Discussion). When *θ*_*i*_ is near 0 or 1, the binomial distribution is highly skewed and larger sample sizes may be required for this approximation to be accurate. Because CentriMo selects the central window from the data, these lower bounds should not be interpreted as exact post-selection confidence bounds with nominal coverage. Instead, we use them as ranking statistics. In our applications, the sample sizes are relatively large and the primary goal is to rank, so we use *L*_*i*_ as the main statistic based on a confidence interval. Nonetheless, we examine *ζ*_*i*_ for comparison.

### Ranking by the lower bound of a Bayesian credible region

As an alternative to the frequentist confidence interval approach, we consider the Bayesian credible region for the parameter *θ*_*i*_ of the binomial distribution, 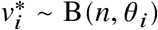 for each motif *i*. We place a Jeffreys prior^28^, Beta(0.5, 0.5), independently on each *θ*_*i*_. Since the beta distribution is a conjugate prior for the binomial distribution^53^, observing *x*_*i*_ successes in *n* trials updates the prior to a posterior *θ*_*i*, posterior_ ∼ Beta (0.5 + *x*_*i*_, 0.5 + *n* − *x*_*i*_). We obtain equal-tailed credible regions through the quantile function of the posterior. For example, for a two-sided 95% equal-tailed credible region, we solve for the 2.5% and the 97.5% quantile of the posterior distribution, to obtain its end points. For the purpose of ranking the motifs, we use the lower endpoint of a two-sided equal-tailed credible region. These credible regions may be asymmetric and are constrained to lie between 0 and 1, unlike the normal-approximation frequentist confidence interval.

Similarly, to rank motifs of width *d*_*i*_ based on effect size, we use the lower endpoint of a credible region, subtracting the expected values under the null hypothesis of no enrichment 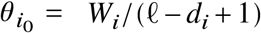. Using the inverse cumulative distribution function 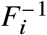 of the posteriors *θ*_*i*, posterior_ ∼ Beta (0.5 + *x*_*i*_, 0.5 + *n* − *x*_*i*_), we will adjust for multiple comparisons with a Bonferroni correction, evaluating at level *α*/2*m*. The correction will ensure that all the credible regions hold simultaneously. Doing this, we obtain

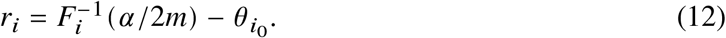

This Bayesian procedure is asymptotically equivalent to using confidence intervals, by the Bernstein– von Mises theorem^61^. Some differences, however, may occur for small samples.

For an empirical-Bayes sensitivity check, we also applied the `ash` function found in AshR^55^ (version 2.2-32), an empirical Bayes approach that shrinks the means of the estimates. The application of the AshR method to our setting requires some special considerations. First, the parameter we are trying to estimate, *θ*, lies in a bounded region between zero and one, so we may have some issues if the parameters lie on the boundary. The credible regions construction as outlined above should not suffer from this problem, however, because the beta distribution used is only defined between 0 and 1, and it is unlikely that the *θ*_0_ is already close to the boundaries. Second, we need to specify the mixture composition to be uniformly positive (`mixcompdist = “+uniform”`), as *θ* always needs to be non-negative.

Having established the four alternative ranking methods and verified the sensitivity of our Bayesian approach, we now evaluate each method’s performance through simulation studies with known ground truth.

### Simulation studies

To evaluate the performance of the different ranking methods, we conducted extensive simulation studies with known ground truth. Without loss of generality, we set *m* = 100 with sample sizes that are either constant or varying across the *m* motifs, to study the impact of sample size *n* on each of the ranking methods.

We constructed four simulation designs to generate the sample sizes *n*. In the respective designs, we:

A. kept *n* constant at 800;
B. drew *n* from U (100, 1500), independent of the true ranking;
C. made *n* proportional to the true ranking (larger *n* for motifs ranked closer to first place); or
D. made *n* inversely proportional to the true ranking.

To simulate the data for each motif *i*, we generated centrally-enriched data by sampling *n* points from a truncated normal distribution, 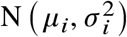, on the interval [−250, 250]. We then ran simulations in two different settings:

a. In Set I, we fix *μ*_*i*_ at 0 and, without loss of generality, randomly draw *σ*_*i*_ from 10 to 120. In this set, motifs with smaller *σ*_*i*_ are more centrally located, so the true ranking is inversely proportional to *σ*_*i*_.
b. In Set II, we fix *σ*_*i*_ at 30 and, without loss of generality, randomly draw *μ*_*i*_ from 0 to 150. In this set, motifs with smaller *μ*_*i*_ are more centrally located, so the true ranking is inversely proportional to *μ*_*i*_.

For each of the eight combinations (2 parameter settings × 4 sample-size designs), we simulated 500 replicates. For each replicate, to best mimic real data, we determined the window size *W*_*i*_ and the corresponding 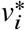 used for the one-sided binomial test using the same method as CentriMo^6^.

We used Kendall’s rank correlation coefficient^17^ to quantify the consistency between the true and sample rankings determined by each method. For each method, we also calculated the average ranking of each motif, across the 500 replicates, for each simulation setting.

## Results

Our application studies included four transcription factor datasets—GATA3 (166 motifs), SRF (123 motifs), ATF1 (262 motifs), and NF-κB-family (92 motifs)—with per-motif sample sizes ranging from 63 to 26 907 (Table 2). Across these applications, the four effect size methods gave broadly similar rankings to one another and, in several cases, ranked the known targeted motif above motifs favoured by p-value ranking.

For each family of motifs under the study, we ranked motifs using the standard p-value approach, and the four alternative effect size methods, namely the modified Cliff’s delta (*δ*^*^; Equation 4), the lower bound of the frequentist asymptotic confidence interval (*L*_*i*_; Equation 8), the lower bound of the frequentist finite-sample confidence interval (*ζ*_*i*_; Equation 11), and the lower bound of the Bayesian credible region (*r*_*i*_; Equation 12).

### Ranking of GATA3-targeted motifs

In humans, the GATA family consists of six transcription factors, named by the common sequence motif they recognize^44^. All six share similar DNA recognition sequences, which makes difficult distinguishing them based on sequence alone. We analysed ChIP-seq data targeting GATA3^64^, attempting to specifically find its motif. In total, we assessed 166 motifs from the JASPAR CORE 2016 vertebrates set^42^, including motifs for the GATA family, supplemented by *de novo* motifs discovered by MEME and DREME from the ChIP-seq data. Notably, the number of sequences containing a motif match (the sample size) *n* ranged from 80 to 5258.

We compared the ranking of the top 10 motifs by CentriMo^6^ p-values and their corresponding ranks determined by the alternative methods (Table 3A). Despite the data examined pertaining to GATA3, CentriMo ranked the GATA2 motif highest. It followed this ranking by a MEME motif generated *de novo* from the ChIP-seq data. CentriMo also ranked GATA1, GATA2, GATA4, and GATA5 above GATA3, which received a ranking of 6. In contrast, all four alternative methods (modified Cliff’s delta, asymptotic and finite-sample confidence intervals, and Bayesian credible region) ranked GATA3 the highest. The four effect size ranking methods all have identical rankings for their top five motifs. We additionally compared this Bayesian credible region ranking against AshR’s empirical-Bayes shrinkage estimate (see Methods); both yielded the same top-5 motifs—GATA3, MEME, GATA2, GATA1, and GATA4 (ranks 1–5).

**Table 3.**
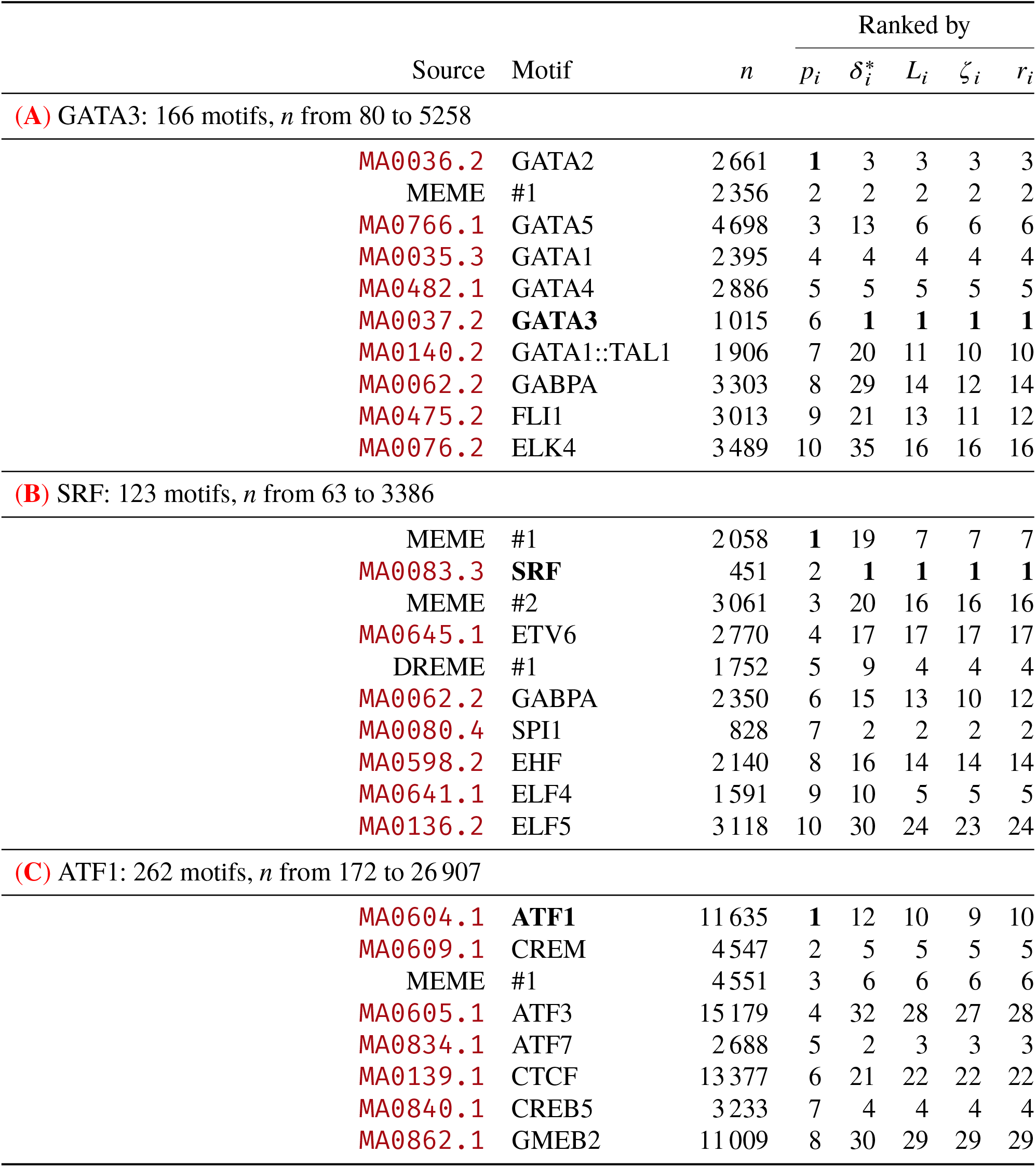
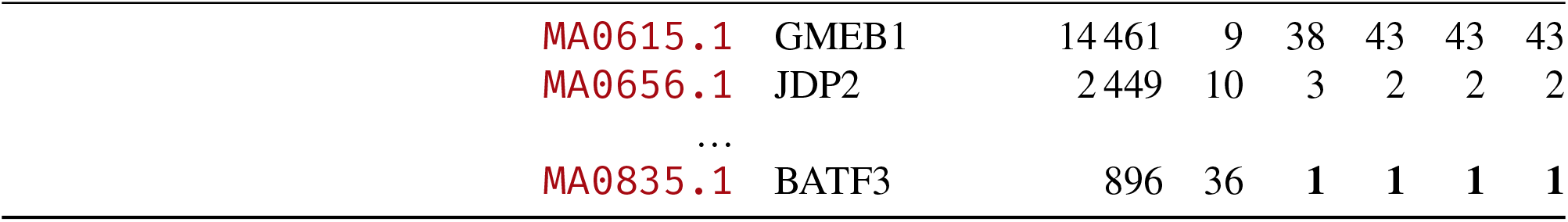
Rankings of peak set-motif correspondences generated by five different methods. Sample sizes and rankings of the top 10 motifs, selected according to their CentriMo^6^ p-values (*p*_*i*_); for ATF1, the 36th-ranked BATF3 motif based on p-value is also shown. The four effect-size methods — modified Cliff’s delta (*δ*^*^; Equation 4), *L*_*i*_ (Equation 8), *ζ*_*i*_ (Equation 11), and *r*_*i*_ (Equation 12) — are defined in Methods. Source: either a JASPAR^42^ ID, or the method used to generate a *de novo* motif from the ChIP-seq data (“MEME”, “DREME”). Motif: either the name of the JASPAR motif, or a number to distinguish between *de novo* from the same source. Bold text motif name: transcription factor pertaining to the ChIP-seq data. Bold text ranking: best rank for some method. Per-motif sample sizes *n* appear in the sub-headers of each panel; total input sequences (peaks) per dataset and other metadata are in Table 2.

The empirical distribution of the best-match positions for each motif reveals that GATA3 has the greatest central enrichment, as compared with GATA5 or GATA2 (Figure 1A). GATA3 (*n* = 1015; *p* = 3.4 × 10^−103^), however, has a sample size nearly five times smaller than GATA5 (*n* = 4698; *p* = 3.0 × 10^−172^). This led to a less significant p-value for GATA3 despite its greater central enrichment. In contrast, the four alternative methods define rankings based on effect size, which is not as sensitive as p-value to sample size. Thus, the effect size methods rank GATA3 at the top position, consistent with the targeted transcription factor, whereas p-value ranking favours motifs with larger sample sizes.

**Figure 1.**
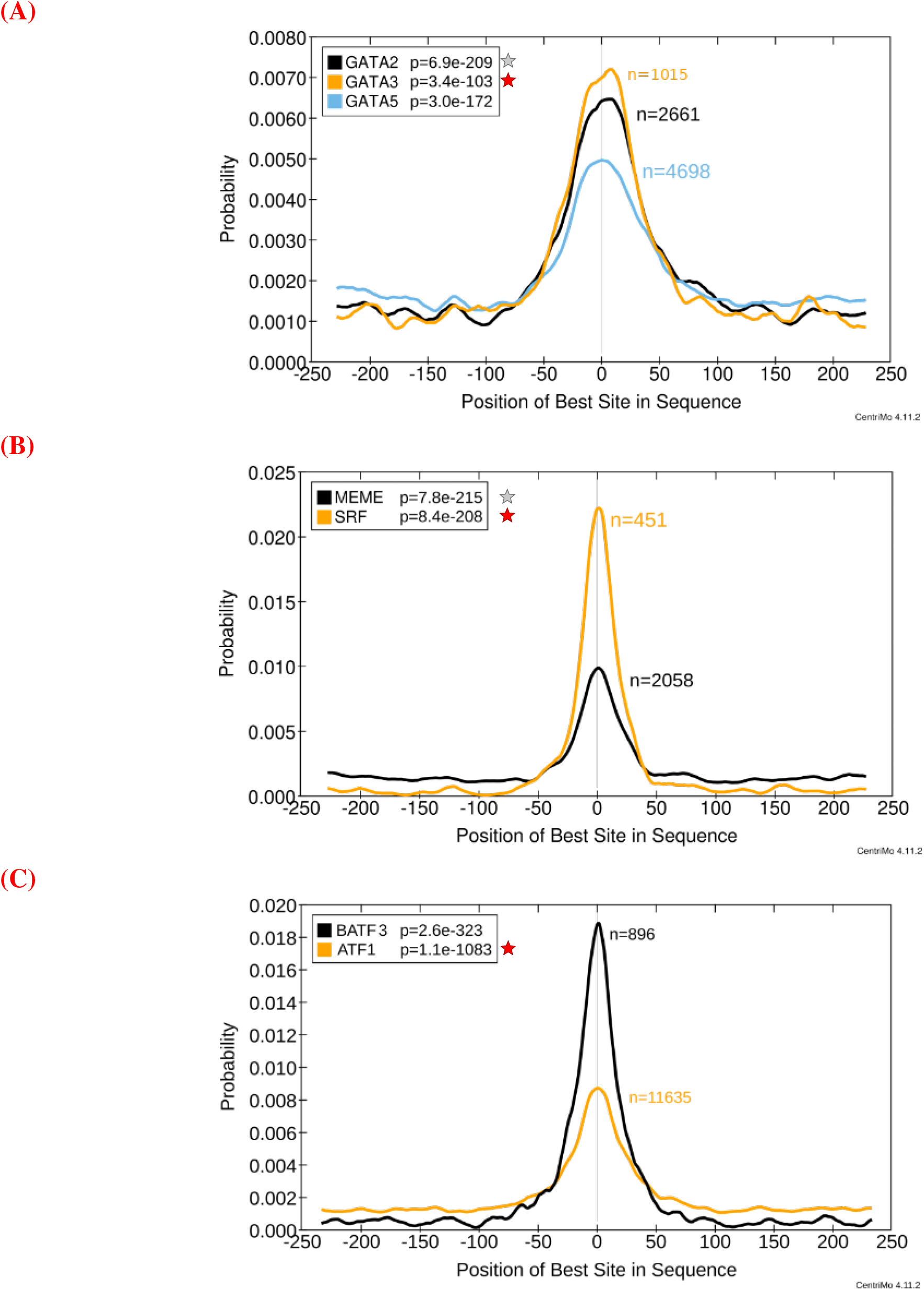
CentriMo^6^probability plots of best motif-match positions within ChIP-seq peaks (smoothing: weighted moving average, window 30). Orange curve with red star (): correct motif; grey star (): the motif with the smallest p-value, which CentriMo accordingly ranks above the correct, more centrally enriched motif (GATA2 in (**A**); the *de novo* MEME motif in (**B**)). MEME: *de novo* motif from ChIP-seq data; legend gives binomial p-values. Experiments target (**A**) GATA3, (**B**) SRF, and (**C**) ATF1.

### Ranking of SRF-targeted motifs

Serum response factor (SRF) regulates cell proliferation and differentiation^47^. Using ChIP-seq data targeting SRF^58^, we applied our five ranking methods. In total, there were 123 motifs, with sample size *n* ranging from 63 to 3386.

Based on p-value, CentriMo^6^ ranked the MEME-generated motif in the first position, above the SRF motif (Table 3B). In terms of central enrichment, however, the SRF motif greatly exceeds the MEME motif (Figure 1B). As with GATA3, the higher p-value ranking of MEME (*n* = 2058) over SRF (*n* = 451) reflects sample size heterogeneity rather than central enrichment magnitude. All four effect size ranking methods place SRF as the top motif, as expected.

### Ranking of ATF1-targeted motifs

We evaluated a CETCh-seq dataset for the cyclic AMP-dependent transcription factor ATF1^52^. Unlike the evaluations above, here, ordering by p-value ranked the targeted motif, ATF1, best (Table 3C). The four alternative methods, however, ranked ATF1 between ninth and 12th. The alternative methods consistently ranked BATF3 in the top position, while ordering by p-value ranked BATF3 only 36th.

Both the ATF1 and BATF3 motifs have highly significant p-values, with p-values < 1 × 10^−300^. BATF3 has a much higher degree of central enrichment than ATF1 (Figure 1C). BATF3, however, has a much smaller sample size (*n* = 896) than ATF1 (*n* = 11 635), a ratio of approximately 13:1. This difference in sample size reduces the p-value rank of BATF3 despite its stronger central enrichment.

The ATF1 analysis illustrates that p-value and effect size rankings can emphasize different aspects of the data. Ranking by p-value reflects both enrichment magnitude and the number of contributing sequences, whereas ranking by effect size more directly reflects the magnitude of central enrichment.

### Ranking of NF-κB-targeted motifs

We analysed the NF-κB family of transcription factors^18^ as a pooled multi-subunit dataset with multiple expected motifs. This distinguishes it from the single-target experiments above. NF-κB proteins associate with each other, regulating inflammation and other immune responses^48^. Using a pooled set of ChIP-seq data of NF-κB subunits (NF-κB1/p50, NF-κB2/p52, RelA/p65, and RelB), CentriMo p-values highly rank the four expected motifs, along with the *de novo* motifs (Table S1); in total 92 motifs were analysed, with sample sizes ranging from 463 to 14 485. Comparable rankings were seen using the alternative effect size ranking methods, with the four motifs of interest and the three generated motifs appearing in the top eight by all five methods (Table S1).

Motifs with larger total numbers of input sequences (that is, larger values of *n*) tend to be ranked higher by p-value than by the effect size methods (Figure 2). Conversely, motifs with smaller numbers of input sequences are ranked relatively lower based on p-values. For each effect size method, we can quantify this by taking the rank by p-value and subtracting rank by the effect size method. This results in a negative Spearman correlation *ρ* between *n* and that difference 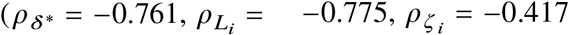, and 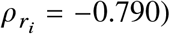.

**Figure 2.**
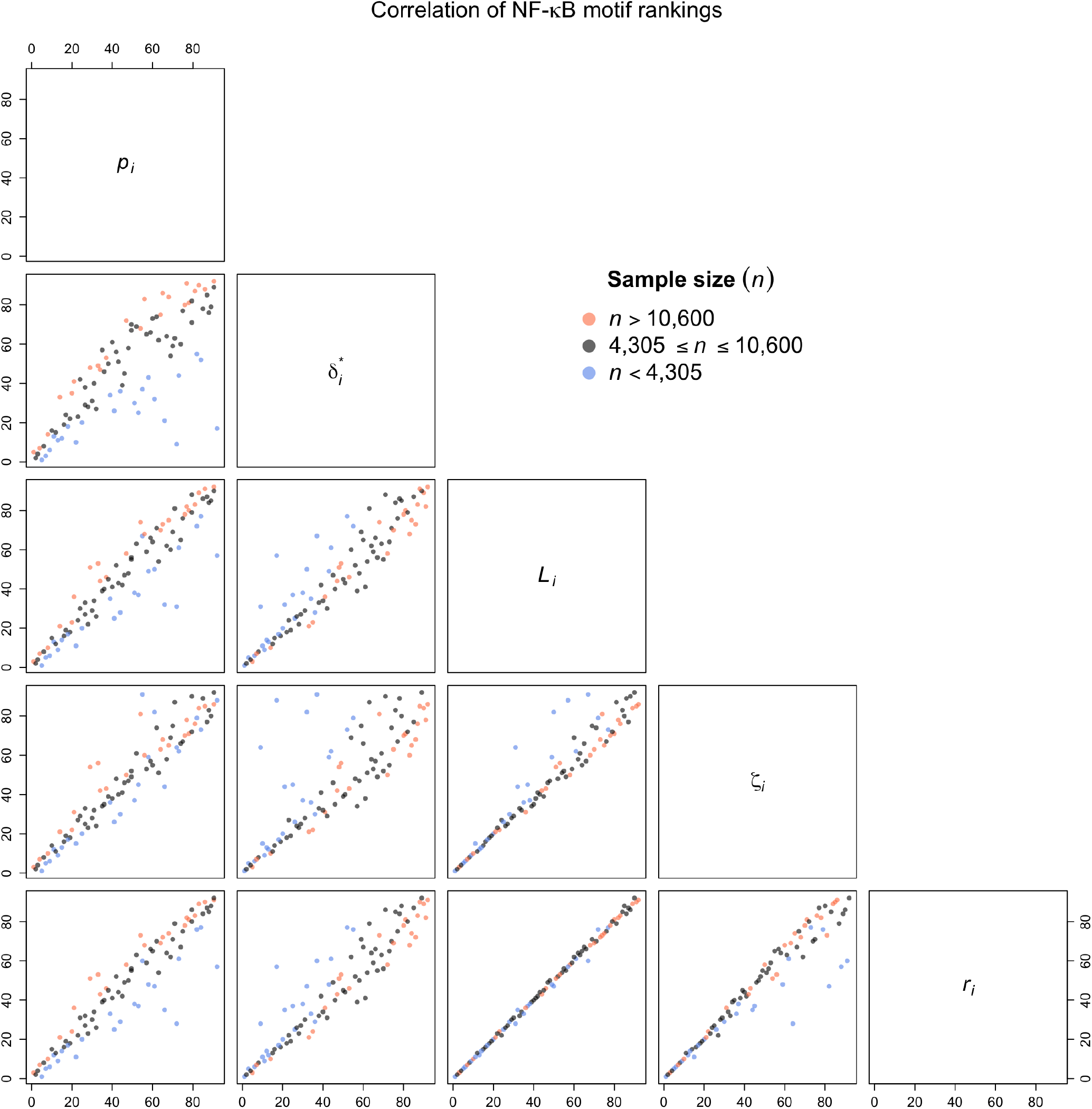
Correlation of NF-κB motif rankings between the CentriMo p-value-method and the four effect size methods (*δ*^*^, *L*_*i*_, *ζ*_*i*_, *r*_*i*_; defined in Methods). Colour: sample size *n* below first quartile (blue), in interquartile range (black), or above third quartile (coral).

### Simulation studies

To further explore the behaviour of our ranking methods, we performed a series of simulation studies. These simulations, both Set I with varying *σ*_*i*_ (Table 4), and Set II with varying *μ*_*i*_ (Table S2), demonstrate the substantial impact that sample size has on ranking by p-value. When the sample sizes did not vary across simulated motifs (Design A), all five methods performed similarly well (Tables 4A, S2A). When we instead randomized the sample sizes independently of the true rankings (Design B), the p-value ranks became highly variable and inaccurate, while the alternative effect size methods remained mostly unaffected (Tables 4B, S2B). Generating the sample sizes to be correlated with the true rankings, pairing the largest sample size with the highest rank (Design C), yielded strong performance by all five methods, as anticipated (Tables 4C, S2C). Finally, pairing the lowest sample size with the highest rank (Design D) caused the p-values to rank in the reverse order, as compared with the true rankings, while the alternative effect size methods remained robust to sample size heterogeneity (Tables 4D, S2D).

**Table 4.** Simulation study results for Set I. True rank: inversely proportional to *σ*_*i*_, that is, motif *i* with the smallest *σ*_*i*_ is most centrally enriched and should be given rank #1. In Set I, *σ*_*i*_ varies and *μ*_*i*_ = 0 for all *i, i* = 1, …, 100. Inferred rank: the mean rank ± standard deviation assigned to a motif with the indicated true rank, averaged across 500 simulated replicates of *m* = 100 motifs. Rankings were determined by p-values (*p*_*i*_), the modified Cliff’s delta (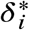; Equation 4), the lower bound of the frequentist asymptotic confidence interval (*L*_*i*_; Equation 8), the lower bound of the frequentist finite-sample confidence interval (*ζ*_*i*_; Equation 11), or the lower bound of the Bayesian credible region (*r*_*i*_; Equation 12). Bold text: the method whose mean rank is closest to the true rank in that row, with ties at displayed precision broken in favour of the smaller standard deviation (the more precise estimate); any methods that remain tied (identical mean rank and standard deviation) are all marked. A perfect method would assign a mean rank equal to the true rank with zero standard deviation. (**A**) Design A: keep *n* constant at 800. (**B**) Design B: draw *n* from U(100, 1500), independently of the true ranking. (**C**) Design C: make *n* proportional to the true ranking. (**D**) Design D: make *n* inversely proportional to the true ranking.

| True rank | Inferred rank |  |  |  |  |
| --- | --- | --- | --- | --- | --- |
| | $p_i$ | $\delta_i^*$ | $L_i$ | $\zeta_i$ | $r_i$ |
| **(A) Constant sample size** |  |  |  |  |  |
| 1 | 1.20 ± 0.49 | **1.13** ± 0.38 | 1.26 ± 0.56 | 1.22 ± 0.51 | 1.26 ± 0.56 |
| 2 | 2.12 ± 0.72 | **2.07** ± 0.55 | 2.13 ± 0.77 | 2.10 ± 0.69 | 2.13 ± 0.77 |
| 3 | **3.00** ± 0.76 | 3.01 ± 0.62 | 3.03 ± 0.91 | 3.01 ± 0.81 | 3.03 ± 0.91 |
| 4 | 4.08 ± 0.86 | **4.00** ± 0.66 | 4.04 ± 0.86 | 4.05 ± 0.81 | 4.04 ± 0.86 |
| 5 | **5.01** ± 0.95 | 5.07 ± 0.77 | 4.95 ± 1.04 | 4.96 ± 0.93 | 4.95 ± 1.04 |
| 6 | 6.08 ± 0.91 | **6.03** ± 0.73 | 6.07 ± 0.91 | 6.07 ± 0.85 | 6.07 ± 0.91 |
| 7 | **7.05** ± 1.05 | 7.08 ± 0.86 | 7.11 ± 1.02 | 7.10 ± 0.95 | 7.11 ± 1.02 |
| 8 | 8.08 ± 1.00 | **7.99** ± 0.89 | 8.06 ± 1.14 | 8.05 ± 1.05 | 8.06 ± 1.14 |
| 9 | **9.04** ± 1.18 | 9.07 ± 0.91 | 9.08 ± 1.17 | 9.06 ± 1.07 | 9.08 ± 1.17 |
| 10 | 10.03 ± 1.13 | **10.00** ± 0.90 | 9.99 ± 1.20 | 9.94 ± 1.07 | 9.99 ± 1.20 |
| **(B) Random sample size** |  |  |  |  |  |
| 1 | 13.26 ± 16.04 | **1.15** ± 0.42 | 1.37 ± 0.70 | 2.57 ± 2.67 | 1.58 ± 1.10 |
| 2 | 14.49 ± 16.95 | **2.05** ± 0.60 | 2.21 ± 1.18 | 3.37 ± 3.08 | 2.43 ± 1.67 |
| 3 | 15.38 ± 16.69 | **3.08** ± 0.73 | 3.11 ± 1.16 | 4.03 ± 3.13 | 3.19 ± 1.61 |
| 4 | 14.41 ± 16.92 | **4.07** ± 0.83 | 4.18 ± 1.35 | 4.65 ± 3.35 | 4.23 ± 1.79 |
| 5 | 14.93 ± 15.90 | **5.08** ± 0.81 | 5.13 ± 1.22 | 5.51 ± 3.08 | 5.16 ± 1.60 |
| 6 | 16.64 ± 17.15 | **6.02** ± 0.95 | 6.06 ± 1.40 | 6.56 ± 3.33 | 6.15 ± 1.85 |
| 7 | 17.21 ± 18.44 | **7.03** ± 0.97 | 7.16 ± 1.70 | 7.55 ± 3.88 | 7.28 ± 2.26 |

**Table 4. Simulation study results for Set I. (continued)**
| True rank | Inferred rank |  |  |  |  |
| --- | --- | --- | --- | --- | --- |
| | $p_i$ | $\delta_i^*$ | $L_i$ | $\zeta_i$ | $r_i$ |
| 8 | 18.38 ± 18.27 | **8.01** ± 1.01 | 7.99 ± 1.63 | 8.48 ± 3.73 | 8.06 ± 2.01 |
| 9 | 18.08 ± 17.75 | **9.09** ± 1.03 | 9.12 ± 1.58 | 9.23 ± 3.57 | 9.10 ± 1.95 |
| 10 | 18.13 ± 17.79 | 9.96 ± 1.05 | 9.92 ± 1.47 | **9.97** ± 3.56 | 9.88 ± 2.02 |
| **(C)** Sample size proportional to the true ranking |  |  |  |  |  |
| 1 | **1.08** ± 0.29 | 1.09 ± 0.31 | 1.16 ± 0.42 | 1.13 ± 0.39 | 1.16 ± 0.42 |
| 2 | **2.07** ± 0.47 | 2.08 ± 0.48 | 2.10 ± 0.62 | 2.10 ± 0.58 | 2.10 ± 0.62 |
| 3 | 2.98 ± 0.53 | **2.99** ± 0.60 | 3.03 ± 0.76 | 2.99 ± 0.67 | 3.02 ± 0.76 |
| 4 | 4.02 ± 0.55 | 4.03 ± 0.58 | **4.00** ± 0.79 | 4.02 ± 0.67 | **4.00** ± 0.79 |
| 5 | 4.98 ± 0.53 | **5.00** ± 0.59 | 5.01 ± 0.73 | 5.00 ± 0.65 | 5.01 ± 0.73 |
| 6 | 6.03 ± 0.54 | **5.99** ± 0.60 | 6.03 ± 0.76 | 6.04 ± 0.69 | 6.03 ± 0.76 |
| 7 | **7.01** ± 0.59 | 7.04 ± 0.66 | 7.02 ± 0.84 | 7.03 ± 0.74 | 7.02 ± 0.84 |
| 8 | **8.01** ± 0.63 | 8.05 ± 0.73 | 8.08 ± 0.95 | 8.05 ± 0.86 | 8.08 ± 0.95 |
| 9 | 9.04 ± 0.59 | 9.01 ± 0.65 | 9.00 ± 0.90 | 9.01 ± 0.81 | **9.00** ± 0.89 |
| 10 | **9.98** ± 0.64 | 10.03 ± 0.78 | 9.98 ± 0.91 | 9.98 ± 0.86 | 9.98 ± 0.90 |
| **(D)** Sample size inversely proportional to the true ranking |  |  |  |  |  |
| 1 | 83.47 ± 9.52 | **1.37** ± 0.68 | 1.97 ± 1.29 | 4.27 ± 3.53 | 2.47 ± 1.88 |
| 2 | 78.24 ± 13.06 | **2.20** ± 0.92 | 2.62 ± 1.59 | 4.33 ± 3.18 | 2.92 ± 1.92 |
| 3 | 73.79 ± 14.43 | **3.07** ± 1.06 | 3.26 ± 1.66 | 4.36 ± 3.28 | 3.46 ± 2.02 |
| 4 | 69.99 ± 16.75 | **4.10** ± 1.18 | 4.24 ± 1.98 | 5.18 ± 3.59 | 4.43 ± 2.38 |
| 5 | 65.39 ± 17.40 | **5.02** ± 1.22 | 5.16 ± 2.06 | 5.52 ± 3.47 | 5.17 ± 2.52 |
| 6 | 61.86 ± 17.74 | **6.00** ± 1.25 | 6.09 ± 2.05 | 6.06 ± 3.48 | 6.03 ± 2.45 |
| 7 | 58.51 ± 17.76 | **6.99** ± 1.22 | 7.01 ± 2.10 | 6.83 ± 3.50 | 6.93 ± 2.48 |
| 8 | 54.40 ± 19.56 | **8.09** ± 1.30 | 7.85 ± 2.06 | 7.55 ± 3.36 | 7.79 ± 2.38 |
| 9 | 51.68 ± 19.26 | **9.05** ± 1.37 | 8.92 ± 2.06 | 8.51 ± 3.57 | 8.87 ± 2.49 |
| 10 | 47.93 ± 19.78 | 10.15 ± 1.36 | **9.95** ± 2.29 | 9.45 ± 3.54 | 9.88 ± 2.65 |

To assess performance of each ranking method over the entirety of each of the two simulation sets and four simulation designs, we used Kendall’s rank correlation *τ* between the average ranking across the 500 simulated replicates for each method and the true rank (Tables 5, S3). Overall, it is evident that sample size has a strong influence on p-value ranking order.

**Table 5.** Kendall’s rank correlation *τ* between the true ranking and each method’s ranking for Set I, with mean ± standard deviation across the same 500 simulated replicates (*m* = 100 motifs) summarised in Table 4. A method recovering the true ranking exactly attains *τ* = 1; *τ* = 0 indicates no rank-order association. Bold: highest mean correlation for each design.

| $P_i$ | $\delta_i^*$ | $L_i$ | $\zeta_i$ | $r_i$ |
| --- | --- | --- | --- | --- |
| **(A)** Constant sample size |  |  |  |  |
| 0.955 ± 0.007 | **0.961** ± 0.007 | 0.957 ± 0.007 | 0.957 ± 0.007 | 0.956 ± 0.007 |
| **(B)** Random sample size |  |  |  |  |
| 0.596 ± 0.040 | **0.954** ± 0.008 | 0.897 ± 0.014 | 0.897 ± 0.014 | 0.925 ± 0.012 |
| **(C)** Sample size proportional to the true ranking |  |  |  |  |
| **0.974** ± 0.004 | 0.951 ± 0.009 | 0.961 ± 0.006 | 0.961 ± 0.006 | 0.956 ± 0.007 |
| **(D)** Sample size inversely proportional to the true ranking |  |  |  |  |
| 0.478 ± 0.136 | **0.963** ± 0.006 | 0.945 ± 0.008 | 0.945 ± 0.008 | 0.950 ± 0.007 |

While p-values had the best performance for Design C (*τ* = 0.974), they had the worst performance in Designs A, B, and D. The p-value ranking method had a low rank correlation for Design D (*τ* = 0.478), where simulation parameters interact with the assumptions inherent in using p-values for ranking most pathologically. Ranking by p-value yielded low rank correlation for Design B (*τ* = 0.596) also. Only the p-value ranking method ever delivered rank correlations below 0.85.

Among the four alternative methods, the modified Cliff’s delta appears to be the least affected by varying sample size in Set I, where it has the best rank correlation in Designs A, B, and D and is the only method exceeding 0.95 across every design. This Set I advantage reflects what varies between the two sets: Set I varies *σ*_*i*_, which alters the concentration of best-match positions about the peak centre—a shape difference that Cliff’s delta, via signed pairwise distance comparisons and without construction of a window, captures directly. Set II fixes *σ*_*i*_ at 30 and varies only *μ*_*i*_, so motif distributions differ in location rather than shape; the parametric methods (*L*_*i*_, *ζ*_*i*_, *r*_*i*_), which directly bound the central-window proportion, then recover the true ranking more accurately than *δ*^*^. Notably, *L*_*i*_ and *ζ*_*i*_ yield identical Kendall’s *τ* values to three decimal places across every design (Tables 5, S3). All four effect size ranking methods, however, are robust to heterogeneous sample sizes and provide similar ranking results.

In practice, motif selection is often treated as an early-retrieval problem: only the top few ranks are inspected, so a method’s accuracy at small ranks matters more than its overall rank correlation. The simulations let us see the different ranking methods’ early-retrieval performance under different scenarios. For example, at true rank 1 in heterogeneous-*n* Designs B and D, *δ*^*^ assigns mean rank better than 4 in both sets, while p-value places the same motif anywhere from rank 13 (Table 4B) to rank 98 (Table S2D).

## Discussion

Sequence motifs found more frequently near the centre of a ChIP-seq peak often represent direct DNA binding by the targeted transcription factor^6^. Although the most commonly used motif ranking method uses p-values, the magnitude of a p-value does not necessarily reveal the true magnitude of central enrichment of a motif. The latter constitutes an effect size, which differs from statistical significance. Lower p-values may be an asset to establishing peak set-motif correspondence, but central enrichment, and therefore effect size, is generally more important.

P-values have a well-known dependence on sample size^50^. In many practical applications in genomics, however, such as genome-wide association studies, one often compares multiple tests among the same or similar sample sizes. This alleviates issues with comparing p-values.

Unfortunately, in ranking transcription factor binding motifs using ChIP-seq data, sample size often varies by several orders of magnitude. This sample size heterogeneity introduces additional complexities that are not sufficiently accounted for by standard statistical measurements of significance. A test with an extremely small effect size can have an arbitrarily small p-value with a sufficiently large number of input sequences (high sample size).

In the GATA3-targeted dataset, sample sizes ranged from 80 to 5258 across the 166 motifs analysed. Despite GATA3’s greater central enrichment than GATA5, apparent in the empirical distribution of best-match positions (Figure 1A), its smaller sample size (*n* = 1015) led to a greater p-value and lower rank as compared with GATA5 (*n* = 4698). All four alternative effect size ranking methods instead appropriately ranked GATA3 in the top position. Similarly, the effect size ranking methods all ranked the known motif best in an analysis targeting SRF.

Ranking by effect size can mitigate the influence of sample size on rank order, but sometimes one should retain sample size as an important consideration. In the motif ranking problem, a larger sample size implies that the motif appears in more ChIP-seq peaks, which is an indication that the motif indeed has biological relevance. For example, in the ATF1-targeted dataset, the expected ATF1 was ranked in the top position based on p-value from CentriMo^6^, while it received a rank of ∼10 by the alternative methods. The empirical distributions of best-match positions (Figure 1C), however, show that ATF1 is clearly less centrally aligned than BATF3—the top motif that was selected by all four alternative ranking methods. Interestingly, despite the small sample size of BATF3 (*n* = 896), the p-value was 2.6 × 10^−323^. The small sample size suggests the greatly significant p-value arises from a large effect size. BATF3’s small p-value, however, posed no competition with *p* = 1.1 × 10^−1083^ for ATF1, driven by its extremely large sample size (*n* = 11 635).

More broadly, disagreement between p-value and effect-size rankings is itself informative: it identifies motifs whose rank is driven primarily by sample size. In the ATF1 case, the discrepancy may reflect a difference in binding mode—direct, high-affinity binding by BATF3 at a relatively small set of sites versus broader, lower-affinity binding by ATF1 across many more peaks. Resolving this hypothesis requires additional biological investigation.

Our four alternative effect size ranking methods were remarkably consistent in both application and simulation studies. Also, although our Bayesian approach might appear distinct, by using the Jeffreys prior, the credible regions constructed have many desirable frequentist properties, including a good coverage rate.

These methods apply to any *in vivo* transcription factor binding assay built on antibodies. This includes the most widely-used assay, ChIP-seq^29,51^, as well as CETCh-seq^52^, as examined in this work. It also includes more recent assays, such as cleavage under targets and release using nuclease (CUT&RUN)^54^, and its extensions. More broadly, however, our methods could apply to any similar problem that has both counts computed in a manner similar to those in CentriMo and problematic sample size heterogeneity.

There exist many alternative methods to improve elucidation of directly binding transcription factor motifs, beyond statistical analyses of central peak distributions. For instance, MASSIF^9^ can improve upon CentriMo outputs, by orthogonally integrating information from DNA-binding domains, and performing meta-analyses of p-values from other enrichment methods. Beyond the scope of our present work, one could improve upon such methods by incorporating the effect size metrics we have considered here. Meta-analyses, for which methods to integrate significance and effect size are already quite well-developed^12,32^, have particular potential for such incorporation.

Beyond other technical biases, due to similarities between target and confounding motifs, correspondences between one peak set and different motifs often share strong correlations^34,66^. A subset of motif sites that the target transcription factor binds may overlap targets for confounding proteins, with similar sequences. This can cause CentriMo^6^ to call matches for multiple overlapping, and possibly off-target, motifs^30^. An off-target motif so confounded may therefore have fewer matches, but still appear predominantly centrally-enriched. CentriMo can, however, often penalize these confounding motifs, since their fewer matches may still be less statistically significant^6^.

Recent versions of CentriMo also now include a new “Concentration” metric. This is effectively an effect size metric, defined as the total probability of all the positions in the central region whose width is the same size as that used to perform smoothing. Indeed, CentriMo now notes^5^ that this may sometimes be more useful than statistical significance, especially for co-factors, which might be more significant, but have lower concentration. This accords with our analyses, for which we were able to leverage our effect size assessments, to distinguish co-factor (and confounding) motifs, from the main targets.

Our methods extend beyond the Concentration metric in several respects. First, our methods account for estimation uncertainty through confidence or credible bounds. Second, they provide simultaneous coverage guarantees across all motifs via Bonferroni adjustment. Third, our effect size methods are grounded in a formal inferential framework with well-characterized statistical properties. Indeed, CentriMo’s own documentation notes that p-value rankings can be confounded by differences in motif information content^6^, an observation consistent with the sample-size dependence we characterize here.

Our alternative methods can sometimes incorrectly prefer the confounding motif, if its effect size is larger. In such instances of this systematic bias, the CentriMo ranking would be preferable. In the event that the technical biases are not so severe as to preclude the identification of the target factor, however, our effect size ranking metrics can often help, as we have shown.

### Confidence interval methods perform similarly

For our frequentist confidence interval approach, we studied both *L*_*i*_ (Equation 8) based on the central limit theorem and *ζ*_*i*_ (Equation 11) based on Hoeffding’s inequality. The Hoeffding-based bound has guaranteed coverage, but it is also more conservative than the asymptotic bound for two reasons. First, *ζ*_*i*_ effectively uses the worst-case variance of the Bernoulli (*θ*) distribution (*σ* =^1^/4 when *θ* =^1^/2), independent of the data. Second, even after substituting the same worst-case variance into *L*_*i*_ (Equation 8), the Hoeffding multiplier 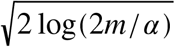 exceeds the asymptotic multiplier *z*_1−*α*/2*m*_ = Φ^−1^(1 − *α*/2*m*) whenever *α*/2*m* ≤ 0.345, which holds for any positive integer *m* as long as *α* ≤ 0.69 (derivation in Supplementary text). This means the asymptotic confidence interval, even using the worst-case variance, is less conservative than the method based on Hoeffding’s inequality, at least for per-family confidence levels greater than 31%. We reiterate, however, that the *ζ*_*i*_ method based on Hoeffding’s inequality has intentional extra conservatism to guarantee sufficient confidence levels in finite samples. By contrast, the interval based on the central limit theorem remains only an asymptotic result. When the sample size grows large enough that the asymptotics “kick in”, the difference between the two confidence interval methods likely stays negligible. We did not observe any practical differences in our previously-described application and simulation studies.

One might also achieve similar results using exact binomial confidence intervals, such as the Clopper–Pearson interval^16,46^. For the sample sizes relevant to motif ranking, the widths of exact and asymptotic intervals are essentially equivalent. Although the asymptotic intervals may have below-nominal coverage, it is the correct ordering of the lower bounds—not exact coverage—that drives motif identification. Small differences in coverage do not produce material differences in lower-bound values or their rank order. A user who wishes to maintain exact coverage can substitute Clopper–Pearson or other exact intervals and expect similar motif identification performance.

### Practical recommendation

For routine ChIP-seq motif analysis with per-motif sample sizes above 100, we recommend the asymptotic confidence interval lower bound (*L*_*i*_; Equation 8). It performs well across both simulation regimes (varying distributional shape in Set I and varying location in Set II), has the simplest closed-form expression, and requires only the counts already computed by CentriMo. The modified Cliff’s delta (*δ*^*^) achieves the highest rank correlation in three of the four Set I designs, where motifs differ in distributional shape, but its computation scales with sample size and it does not provide an explicit confidence bound. The Hoeffding-based bound (*ζ*_*i*_) offers finite-sample coverage guarantees and is preferable when some motifs have very few sequences (fewer than 30). The Bayesian credible lower bound (*r*_*i*_) provides an alternative interpretation but is asymptotically equivalent to *L*_*i*_ by the Bernstein–von Mises theorem.

P-values dominating when sample sizes correlate with the true ranking and effect sizes dominating when they do not motivates the development of an integrated metric. Although p-values themselves implicitly combine effect size and sample size, our simulations show that this combination weights sample size strongly enough to invert the rank order when sample sizes correlate negatively with the truth (Design D), so a deliberately balanced metric could outperform either alternative. Many experiments, however, inherently preclude finding the target motif, due to confounders or simply lack of sufficient quality data. Given that, an integrated metric might not always yield an improvement. Alternatively, such a direction could be considered in future work, as an indicator of a likelihood of technical biases overwhelming the true signal in an experiment. The various effect size metrics we present here, and our simulation studies, are an important first step, towards undertaking these kinds of refinements and extensions.

In practice, we suggest the following workflow: (1) run CentriMo in the usual manner to obtain per-motif central enrichment statistics; (2) apply the R code available from our repository (see Availability) to compute effect-size rankings; (3) rank motifs by the asymptotic confidence interval lower bound *L*_*i*_; and (4) compare with the original p-value rankings. Discrepancies between the two rankings identify motifs whose rank is influenced by sample size rather than enrichment magnitude and may warrant closer inspection.

## Table of notation

| Symbol | Class | Description |
| --- | --- | --- |
| $\alpha$ | Probability | Type I error rate |
| $B$ | Distribution | Binomial |
| $B_{i,j}$ | Random variable | Bernoulli trial indicator; $B_{i,j} \stackrel{\text{iid}}{\sim} \text{Bernoulli}(\theta_i)$ |
| $d$ | Length | ... of motif |
| $\Delta_i$ | Effect size | Departure of $\theta_i$ from its null value; $= \theta_i - \theta_{i_0}$ |
| $\delta$ | Effect size | Cliff's delta |
| $\delta^*$ | Effect size | Cliff's delta for magnitudes |
| $\epsilon$ | Bound | Deviation bound in Hoeffding's inequality |
| $H$ | Hypothesis | |
| $H_0$ | Hypothesis | Null |
| $i$ | Motif | |
| $j$ | Position | |
| $\ell$ | Length | ... of input sequences |
| $\ell_i$ | Length | ... of valid starting-position range for motif $i$, $= \ell - d_i + 1$ |
| $L_i$ | Effect size | Asymptotic CI lower bound (Equation 8) |
| $\mu$ | Number | Mean |
| $m$ | Number | ... of motifs |
| $n$ | Number | ... of input sequences |
| $P$ | Probability | |
| $p$ | P-value | (Equation 1) |
| $p_i$ | P-value | ... for motif $i$ |
| $\Phi$ | Function | Cumulative distribution function of standard normal distribution |
| $r_i$ | Effect size | Bayesian credible lower bound |
| $\sigma$ | Number | Population standard deviation |
| $\theta$ | Probability | ... of success for each trial |
| $\theta_0$ | Probability | ... of success for each trial under $H_0$, $= W/(\ell - d + 1)$ |
| $\theta_{i_0}$ | Probability | ... of success for each trial under $H_0$, $= W_i/(\ell - d_i + 1)$ |
| $U$ | Distribution | Uniform |
| $U$ | Interval | Reference uniform support over $[-\ell_i/2, \ell_i/2]$ |
| $V_i$ | Multiset | Positions of best matches of motif $i$ |
| $v_{i,j}$ | Number | Multiplicity of best matches of motif $i$ starting at position $j$ |
| $v_i^*$ | Number | Multiplicity of best matches of motif $i$ starting at positions in central window |
| $W$ | Length | ... of central window, $W < \ell$ |
| $X$ | Set | |
| $x$ | Element | ... of $X$ |
| $Y$ | Set | |
| $y$ | Element | ... of $Y$ |
| $z$ | Quantile | Standard normal quantile; $z_p = \Phi^{-1}(p)$ |
| $\zeta_i$ | Effect size | Finite-sample CI lower bound |

## Authors’ contributions

Conceptualization, C.V. and M.M.H.; Data Curation, C.V. and D.D.; Formal Analysis, S.M., J.N., Y.T., L.Z., and L.S.; Investigation, S.M., C.V., and L.S.; Methodology, S.M., C.V., D.D., J.N., Y.T., L.Z., M.M.H., and L.S.; Software, S.M. and C.V.; Visualization, S.M.; Writing — Original Draft, S.M., C.V., Y.T., L.Z., and L.S.; Writing — Review & Editing, S.M., C.V., J.N., M.M.H., and L.S.; Resources, M.M.H. and L.S.; Funding Acquisition, M.M.H. and L.S.; Project Administration, C.V., M.M.H., and L.S.; Supervision, M.M.H. and L.S. Middle authors (D.D., J.N., Y.T., and L.Z.) are listed alphabetically.

## Acknowledgments

We thank Christopher K. Glass and Verena Link for providing SRF datasets. We thank Carl Virtanen and Zhibin Lu (Bioinformatics and High Performance Computing Core, University Health Network), for technical assistance.

## Availability

We provide R source code for our motif rankings (https://github.com/ScottMastro/motif-ranking). Persistent availability is ensured by Zenodo, in which we have deposited the version of our code we used (https://doi.org/10.5281/zenodo.20727949) and the CentriMo outputs we utilized (https://doi.org/10.5281/zenodo.20728417). All of our source code is licensed under a GNU General Public License, version 3 (GPLv3).

## Supplementary tables

**Table S1.** Rankings of peak set-motif correspondence for NF-κB-targeted motifs generated by five different methods. This analysis included 92 motifs, with sample sizes *n* ranging from 463 to 14 485. The top 10 motifs, selected according to their CentriMo^6^ p-values (*p*), are shown. The NF-κB dataset uses pooled ChIP-seq data from four NF-κB subunits (NF-κB1/p50, NF-κB2/p52, RelA/p65, and RelB)^18^. Motifs labelled “MEME” or “DREME” indicate a *de novo* motif ranked at the indicated position. Rankings were also computed using the modified Cliff’s delta (*δ*^*^; Equation 4), the lower bound of the frequentist asymptotic confidence interval (*L*_*i*_; Equation 8), the lower bound of the frequentist finite-sample confidence interval (*ζ*_*i*_; Equation 11), and the lower bound of the Bayesian credible region (*r*_*i*_; Equation 12). Source: either a JASPAR^42^ ID, or the method used to generate a *de novo* motif from the ChIP-seq data (“MEME”, “DREME”). Motif: either the name of the JASPAR motif, or a number to distinguish between *de novo* from the same source. Bold text motif name: the 4 transcription factors pertaining to the ChIP-seq data. Bold text ranking: top 4 ranks for some method.

|  |  |  |  | Ranked by |  |  |  |  |
| --- | --- | --- | --- | --- | --- | --- | --- | --- |
| Source | Motif | $n$ | $p$ | $\delta^*$ | $L_i$ | $\zeta_i$ | $r_i$ | |
| NF-κB, 92 motifs, $n$ from 463 to 14 485 | | | | | | | | |
| MEME | #1 | 12 406 | **1** | 5 | **3** | **3** | **3** |  |
| MA0778.1 | **NFKB2** | 8 295 | **2** | **2** | **2** | **2** | **2** |  |
| MA0107.1 | **RELA** | 9 944 | **3** | **4** | **4** | **4** | **4** |  |
| MA0101.1 | **REL** | 13 475 | **4** | 7 | 7 | 7 | 7 |  |
| MA0105.4 | **NFKB1** | 3 858 | 5 | **1** | **1** | **1** | **1** |  |
| DREME | #1 | 10 081 | 6 | 8 | 8 | 8 | 8 |  |
| DREME | #2 | 2 721 | 7 | **3** | 5 | 5 | 5 |  |
| MA0056.1 | MZF1 | 14 485 | 8 | 14 | 10 | 10 | 10 |  |
| DREME | #3 | 2 869 | 9 | 6 | 6 | 6 | 6 |  |
| DREME | #4 | 9 317 | 10 | 16 | 15 | 14 | 15 |  |

**Table S2.**
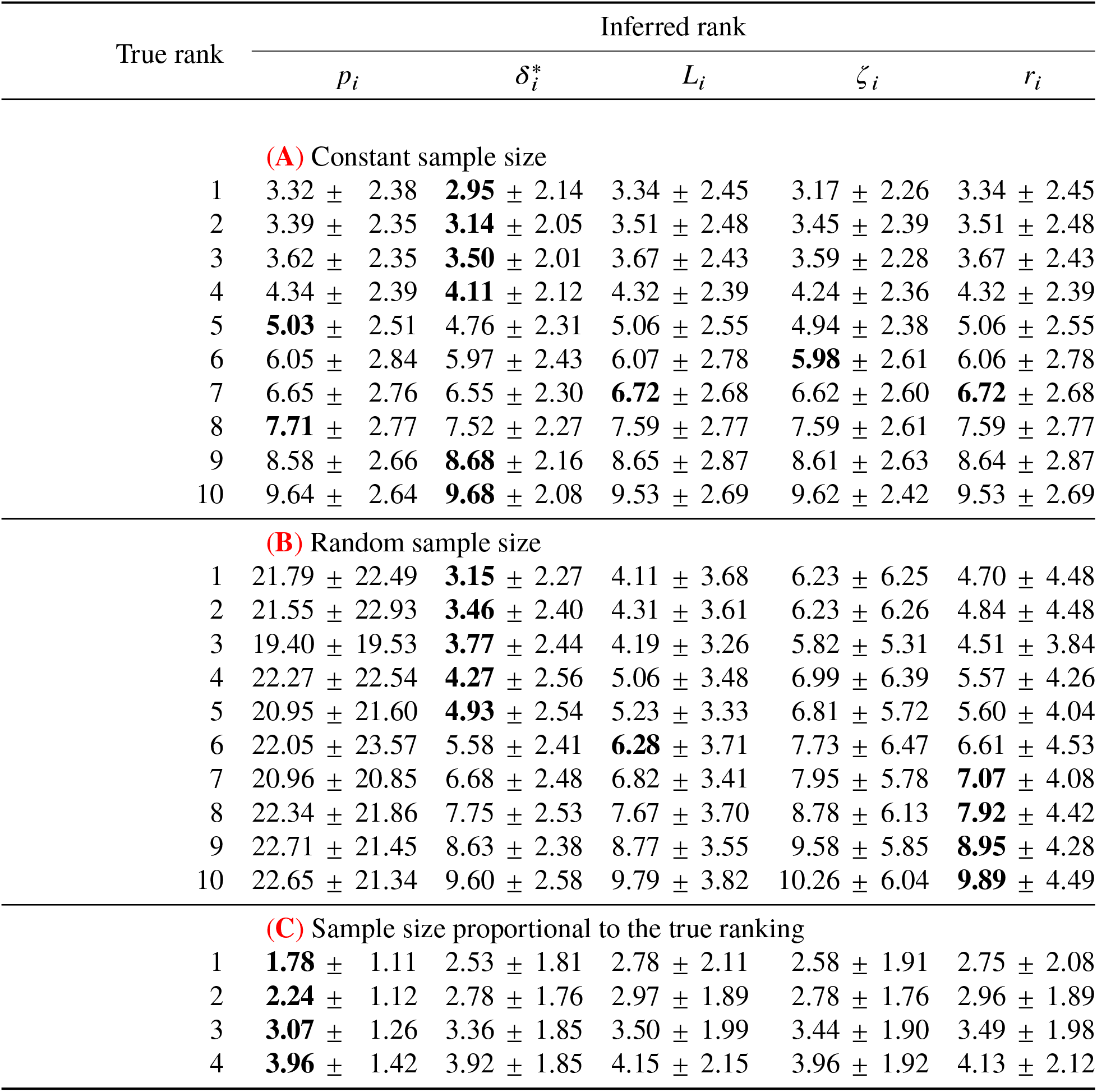
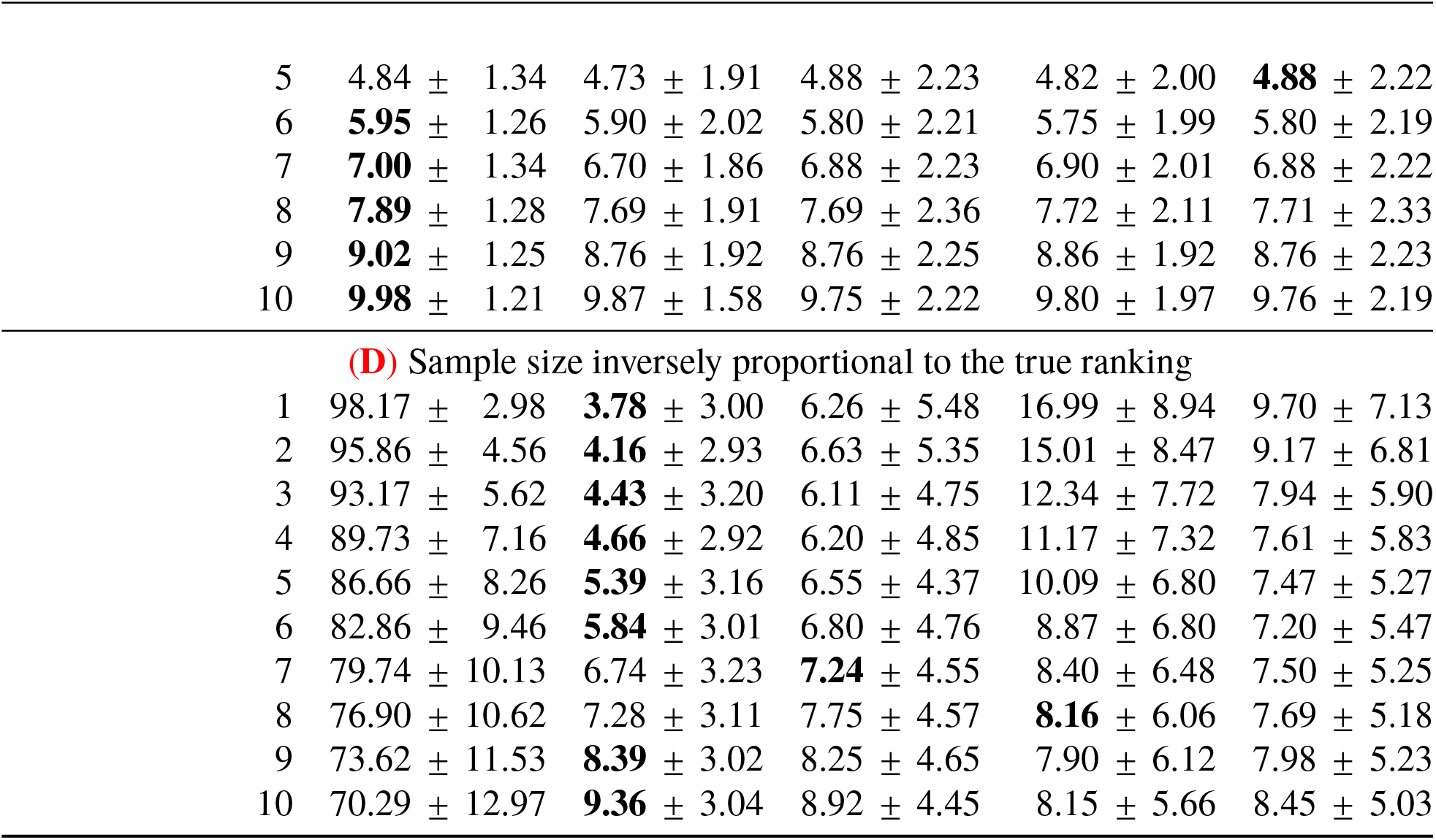
Simulation study results for Set II. True rank: inversely proportional to *μ*_*i*_, that is, motif *i* with the smallest *μ*_*i*_ is most centrally enriched and should be given rank #1. In Set II, *σ*_*i*_ = 30 for all *i*, and *μ*_*i*_ varies for *i* = 1, …, 100. Inferred rank: the mean rank ± standard deviation assigned to a motif with the indicated true rank, averaged across 500 simulated replicates of *m* = 100 motifs. Rankings were determined by p-values (*p*_*i*_), the modified Cliff’s delta (*δ*^*^; Equation 4), the lower bound of the frequentist asymptotic confidence interval (*L*_*i*_; Equation 8), the lower bound of the frequentist finite-sample confidence interval (*ζ*_*i*_; Equation 11), or the lower bound of the Bayesian credible region (*r*_*i*_; Equation 12). Bold text: the method whose mean rank is closest to the true rank in that row, with ties at displayed precision broken in favour of the smaller standard deviation (the more precise estimate); any methods that remain tied (identical mean rank and standard deviation) are all marked. A perfect method would assign a mean rank equal to the true rank with zero standard deviation. (A) Design A: keep *n* constant at 800. (**B**) Design B: draw *n* from U(100, 1500), independently of the true ranking. (**C**) Design C: make *n* proportional to the true ranking. (**D**) Design D: make *n* inversely proportional to the true ranking.

| True rank | Inferred rank |  |  |  |  |
| --- | --- | --- | --- | --- | --- |
| | $p_i$ | $\delta_i^*$ | $L_i$ | $\zeta_i$ | $r_i$ |
| **(A)** Constant sample size |  |  |  |  |  |
| 1 | 3.32 ± 2.38 | **2.95** ± 2.14 | 3.34 ± 2.45 | 3.17 ± 2.26 | 3.34 ± 2.45 |
| 2 | 3.39 ± 2.35 | **3.14** ± 2.05 | 3.51 ± 2.48 | 3.45 ± 2.39 | 3.51 ± 2.48 |
| 3 | 3.62 ± 2.35 | **3.50** ± 2.01 | 3.67 ± 2.43 | 3.59 ± 2.28 | 3.67 ± 2.43 |
| 4 | 4.34 ± 2.39 | **4.11** ± 2.12 | 4.32 ± 2.39 | 4.24 ± 2.36 | 4.32 ± 2.39 |
| 5 | **5.03** ± 2.51 | 4.76 ± 2.31 | 5.06 ± 2.55 | 4.94 ± 2.38 | 5.06 ± 2.55 |
| 6 | 6.05 ± 2.84 | 5.97 ± 2.43 | 6.07 ± 2.78 | **5.98** ± 2.61 | 6.06 ± 2.78 |
| 7 | 6.65 ± 2.76 | 6.55 ± 2.30 | **6.72** ± 2.68 | 6.62 ± 2.60 | **6.72** ± 2.68 |
| 8 | **7.71** ± 2.77 | 7.52 ± 2.27 | 7.59 ± 2.77 | 7.59 ± 2.61 | 7.59 ± 2.77 |
| 9 | 8.58 ± 2.66 | **8.68** ± 2.16 | 8.65 ± 2.87 | 8.61 ± 2.63 | 8.64 ± 2.87 |
| 10 | 9.64 ± 2.64 | **9.68** ± 2.08 | 9.53 ± 2.69 | 9.62 ± 2.42 | 9.53 ± 2.69 |
| **(B)** Random sample size |  |  |  |  |  |
| 1 | 21.79 ± 22.49 | **3.15** ± 2.27 | 4.11 ± 3.68 | 6.23 ± 6.25 | 4.70 ± 4.48 |
| 2 | 21.55 ± 22.93 | **3.46** ± 2.40 | 4.31 ± 3.61 | 6.23 ± 6.26 | 4.84 ± 4.48 |
| 3 | 19.40 ± 19.53 | **3.77** ± 2.44 | 4.19 ± 3.26 | 5.82 ± 5.31 | 4.51 ± 3.84 |
| 4 | 22.27 ± 22.54 | **4.27** ± 2.56 | 5.06 ± 3.48 | 6.99 ± 6.39 | 5.57 ± 4.26 |
| 5 | 20.95 ± 21.60 | **4.93** ± 2.54 | 5.23 ± 3.33 | 6.81 ± 5.72 | 5.60 ± 4.04 |
| 6 | 22.05 ± 23.57 | 5.58 ± 2.41 | **6.28** ± 3.71 | 7.73 ± 6.47 | 6.61 ± 4.53 |
| 7 | 20.96 ± 20.85 | 6.68 ± 2.48 | 6.82 ± 3.41 | 7.95 ± 5.78 | **7.07** ± 4.08 |
| 8 | 22.34 ± 21.86 | 7.75 ± 2.53 | 7.67 ± 3.70 | 8.78 ± 6.13 | **7.92** ± 4.42 |
| 9 | 22.71 ± 21.45 | 8.63 ± 2.38 | 8.77 ± 3.55 | 9.58 ± 5.85 | **8.95** ± 4.28 |
| 10 | 22.65 ± 21.34 | 9.60 ± 2.58 | 9.79 ± 3.82 | 10.26 ± 6.04 | **9.89** ± 4.49 |
| **(C)** Sample size proportional to the true ranking |  |  |  |  |  |
| 1 | **1.78** ± 1.11 | 2.53 ± 1.81 | 2.78 ± 2.11 | 2.58 ± 1.91 | 2.75 ± 2.08 |
| 2 | **2.24** ± 1.12 | 2.78 ± 1.76 | 2.97 ± 1.89 | 2.78 ± 1.76 | 2.96 ± 1.89 |
| 3 | **3.07** ± 1.26 | 3.36 ± 1.85 | 3.50 ± 1.99 | 3.44 ± 1.90 | 3.49 ± 1.98 |
| 4 | **3.96** ± 1.42 | 3.92 ± 1.85 | 4.15 ± 2.15 | 3.96 ± 1.92 | 4.13 ± 2.12 |

**Table S2.** Simulation study results for Set II. (*continued*)
| True rank | Inferred rank |  |  |  |  |
| --- | --- | --- | --- | --- | --- |
| | $p_i$ | $\delta_i^*$ | $L_i$ | $\zeta_i$ | $r_i$ |
| 5 | 4.84 ± 1.34 | 4.73 ± 1.91 | 4.88 ± 2.23 | 4.82 ± 2.00 | **4.88** ± 2.22 |
| 6 | **5.95** ± 1.26 | 5.90 ± 2.02 | 5.80 ± 2.21 | 5.75 ± 1.99 | 5.80 ± 2.19 |
| 7 | **7.00** ± 1.34 | 6.70 ± 1.86 | 6.88 ± 2.23 | 6.90 ± 2.01 | 6.88 ± 2.22 |
| 8 | **7.89** ± 1.28 | 7.69 ± 1.91 | 7.69 ± 2.36 | 7.72 ± 2.11 | 7.71 ± 2.33 |
| 9 | **9.02** ± 1.25 | 8.76 ± 1.92 | 8.76 ± 2.25 | 8.86 ± 1.92 | 8.76 ± 2.23 |
| 10 | **9.98** ± 1.21 | 9.87 ± 1.58 | 9.75 ± 2.22 | 9.80 ± 1.97 | 9.76 ± 2.19 |
| **(D)** Sample size inversely proportional to the true ranking |  |  |  |  |  |
| 1 | 98.17 ± 2.98 | **3.78** ± 3.00 | 6.26 ± 5.48 | 16.99 ± 8.94 | 9.70 ± 7.13 |
| 2 | 95.86 ± 4.56 | **4.16** ± 2.93 | 6.63 ± 5.35 | 15.01 ± 8.47 | 9.17 ± 6.81 |
| 3 | 93.17 ± 5.62 | **4.43** ± 3.20 | 6.11 ± 4.75 | 12.34 ± 7.72 | 7.94 ± 5.90 |
| 4 | 89.73 ± 7.16 | **4.66** ± 2.92 | 6.20 ± 4.85 | 11.17 ± 7.32 | 7.61 ± 5.83 |
| 5 | 86.66 ± 8.26 | **5.39** ± 3.16 | 6.55 ± 4.37 | 10.09 ± 6.80 | 7.47 ± 5.27 |
| 6 | 82.86 ± 9.46 | **5.84** ± 3.01 | 6.80 ± 4.76 | 8.87 ± 6.80 | 7.20 ± 5.47 |
| 7 | 79.74 ± 10.13 | 6.74 ± 3.23 | **7.24** ± 4.55 | 8.40 ± 6.48 | 7.50 ± 5.25 |
| 8 | 76.90 ± 10.62 | 7.28 ± 3.11 | 7.75 ± 4.57 | **8.16** ± 6.06 | 7.69 ± 5.18 |
| 9 | 73.62 ± 11.53 | **8.39** ± 3.02 | 8.25 ± 4.65 | 7.90 ± 6.12 | 7.98 ± 5.23 |
| 10 | 70.29 ± 12.97 | **9.36** ± 3.04 | 8.92 ± 4.45 | 8.15 ± 5.66 | 8.45 ± 5.03 |

**Table S3.** Kendall’s rank correlation *τ* between the true ranking and each method’s ranking for Set II, with mean ± standard deviation across the same 500 simulated replicates (*m* = 100 motifs) summarised in Table S2. A method recovering the true ranking exactly attains *τ* = 1; *τ* = 0 indicates no rank-order association. Bold: highest mean correlation for each design.

| $p_i$ | $\delta_i^*$ | $L_i$ | $\zeta_i$ | $r_i$ |
| --- | --- | --- | --- | --- |
| **(A)** Constant sample size |  |  |  |  |
| 0.952 ± 0.006 | 0.877 ± 0.035 | **0.968** ± 0.004 | **0.968** ± 0.004 | 0.963 ± 0.005 |
| **(B)** Random sample size |  |  |  |  |
| 0.541 ± 0.045 | 0.872 ± 0.036 | 0.894 ± 0.012 | 0.894 ± 0.012 | **0.933** ± 0.009 |
| **(C)** Sample size proportional to the true ranking |  |  |  |  |
| **0.973** ± 0.004 | 0.877 ± 0.036 | **0.973** ± 0.004 | **0.973** ± 0.004 | 0.964 ± 0.005 |
| **(D)** Sample size inversely proportional to the true ranking |  |  |  |  |
| 0.187 ± 0.152 | 0.869 ± 0.038 | 0.931 ± 0.017 | 0.931 ± 0.017 | **0.944** ± 0.010 |

## Supplementary text

### Comparison of confidence interval methods

By factoring Equation 11 as below, we can better clarify the two sources of *ζ*_*i*_’s extra conservatism relative to *L*_*i*_,

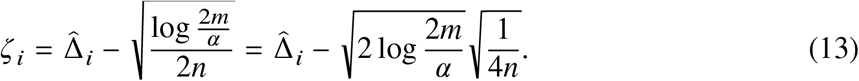

Firstly, *ζ*_*i*_ effectively uses the worst-case variance of the Bernoulli (*θ*) distribution (*σ* =^1^/4 when *θ* =^1^/2) to compute the confidence interval. Since the true parameter value is unknown, using the worst-case variance is necessary to guarantee coverage, irrespective of the true effect size.

If we also assume the worst-case variance for *L*_*i*_ (Equation 8), then we get a bound that is more conservative than *L*_*i*_ itself, since

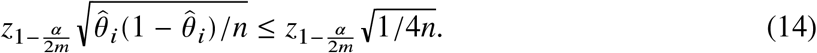

Thus, to finish reconciling the differences between *L*_*i*_ (Equation 8) and *ζ*_*i*_ (Equation 11) we need only compare the multipliers of 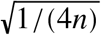 (Equation 13 and Equation 14), since the rest of the bounds are identical. Namely, we compare

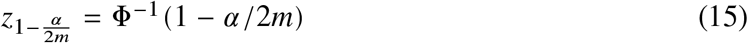

with

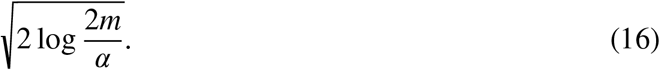

To this end, we note that

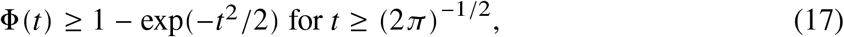

or equivalently

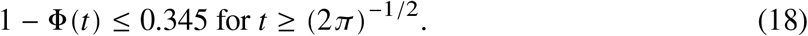

Therefore,

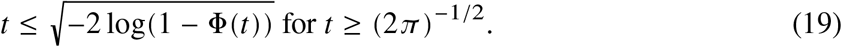

Furthermore,

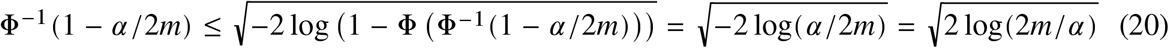

whenever *α*/2*m* ≤ 0.345. This holds for any positive integer *m*, as long as *α* ≤ 0.69.

Thus,

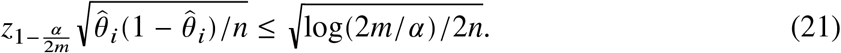

This confirms the claim: the asymptotic confidence interval, even using the worst-case variance, is less conservative than the method based on Hoeffding’s inequality for any per-family confidence level above 31%.

